# A novel dimerization module in Gemin5 is critical for protein recruitment and translation control

**DOI:** 10.1101/654111

**Authors:** María Moreno-Morcillo, Rosario Francisco-Velilla, Azman Embarc-Buh, Javier Fernández-Chamorro, Santiago Ramón-Maiques, Encarnación Martínez-Salas

## Abstract

The versatile multifunctional protein Gemin5 is involved in small nuclear ribonucleoproteins (snRNPs) assembly, ribosome binding, and translation control through distinct domains located at the protein ends. However, the structure and function of the central moiety of Gemin5 remained unknown. Here, we solved the crystal structure of an extended tetratricopeptide (TPR)-like domain in the middle region of Gemin5, demonstrating that it self-assembles into a canoe-shaped dimer. Mass spectrometry analysis shows that this dimerization module is functional in living cells and drives the interaction between p85, a viral-induced Gemin5 cleavage fragment, and the full-length Gemin5. In contrast, disruption of the dimerization surface by a point mutation in the TPR-like domain prevents this interaction and abrogates the translation enhancement induced by p85. The structural characterization of this unprecedented dimerization domain provides the mechanistic basis for a role of the middle region of Gemin5 as a key mediator of protein-protein interactions.

**HIGHLIGHTS:**

- The crystal structure of a central region of Gemin5 reveals a novel dimerization domain
- The proteolytic product of Gemin5 (p85) recruits the endogenous protein through the dimerization module
- The dimerization capability of Gemin5 determines the factors recruited in human cells
- Disruption of the dimerization domain impairs p85 ability to stimulate translation

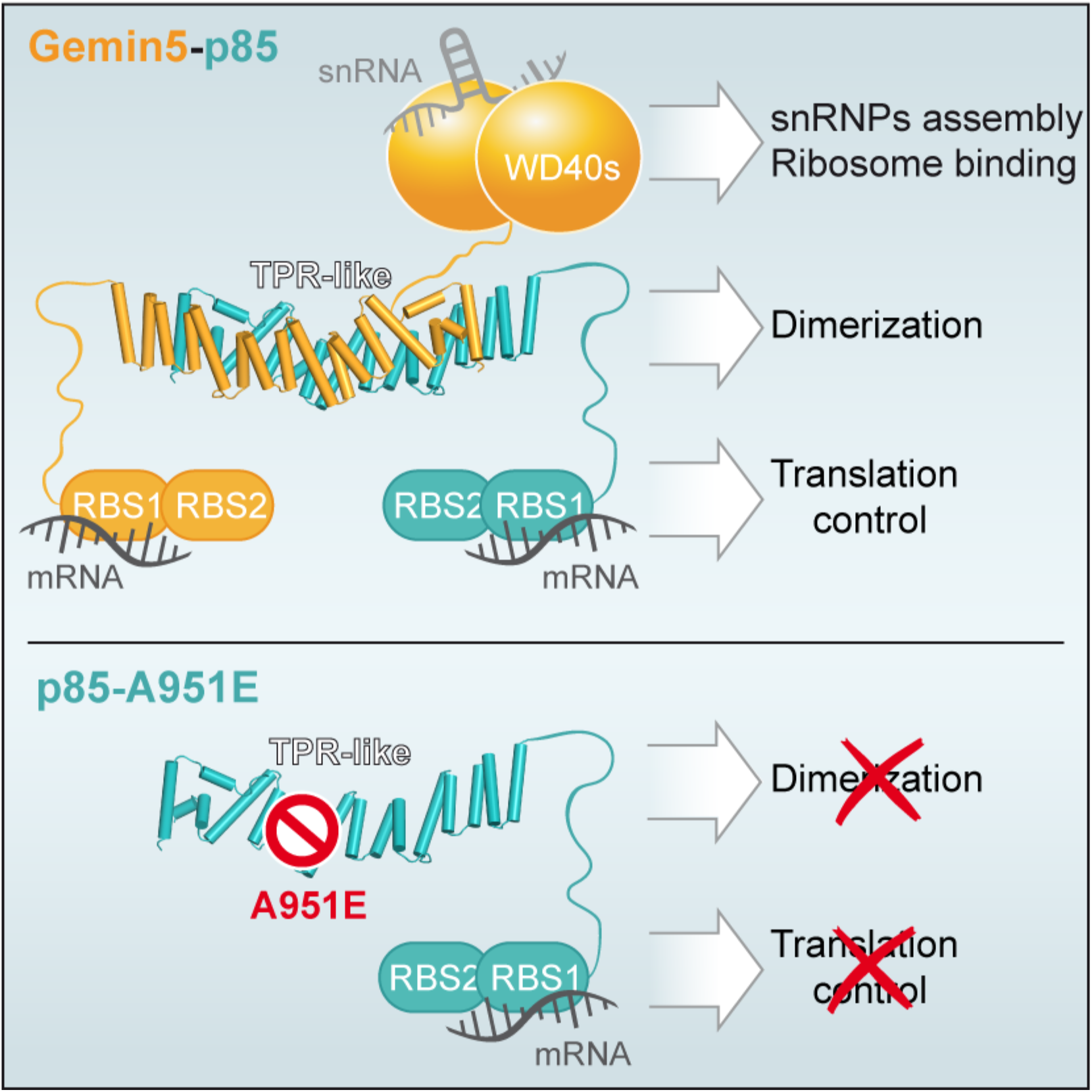
Graphical abstract.

## INTRODUCTION

RNA-binding proteins (RBPs) play a pivotal role in gene expression control and cell homeostasis (Gehring et al., 2017). Generally, RBPs comprise RNA-binding domains (RBD) and protein-protein interaction modules (Ellisdon et al., 2012; Katibah et al., 2013; Lunde et al., 2007; Yang et al., 2012), such that the combination of distinct domains provides multiple features to these factors. Gemin5 is a predominantly cytoplasmic RBP that forms part of the survival of motor neuron (SMN) complex in metazoan organisms (Battle et al., 2006; Matera et al., 2019). This multi-protein complex plays a critical role in the biogenesis of small nuclear ribonucleoproteins (snRNPs) (Lau et al., 2009), the components of the splicing machinery. However, Gemin5 is mainly found outside of the SMN complex (Battle et al., 2007), suggesting that it may have additional functions. In agreement with this view, Gemin5 acts as a scaffold protein, serving as a hub for distinct ribonucleoprotein (RNP) networks. Indeed, Gemin5 has been identified as a down-regulator of translation (Francisco-Velilla et al., 2018; Pacheco et al., 2009; Workman et al., 2015), and as a ribosome-interacting factor (Francisco-Velilla et al., 2016; Simsek et al., 2017).

RBPs perform critical functions in all organisms, including viruses. However, viruses are obliged pathogens with reduced coding capacity and thus, have developed various strategies to subvert essential host factors into their own benefit, which include the proteolysis of specific RBPs and initiation factors (eIFs) (Walsh and Mohr, 2011). In particular, RNA viruses exemplified by picornaviruses, modify host factors to promote translation of the viral RNA using cap-independent mechanisms governed by IRES (Internal Ribosome Entry Site) elements (Lozano and Martinez-Salas, 2015), evading the inhibition of cap-dependent translation occurring in infected cells. Consistent with its role in key cellular processes, Gemin5 is proteolytically cleaved in picornavirus-infected cells, producing a polypeptide of 85 kDa (thereafter p85) (Figure 1A) (Pineiro et al., 2012). Importantly, contrary to the negative effect of Gemin5 in translation (Pacheco et al., 2009), expression of p85 in human cells stimulates IRES-driven translation (Fernandez-Chamorro et al., 2014). The basis of this different behavior has been poorly studied.

**Figure 1.**
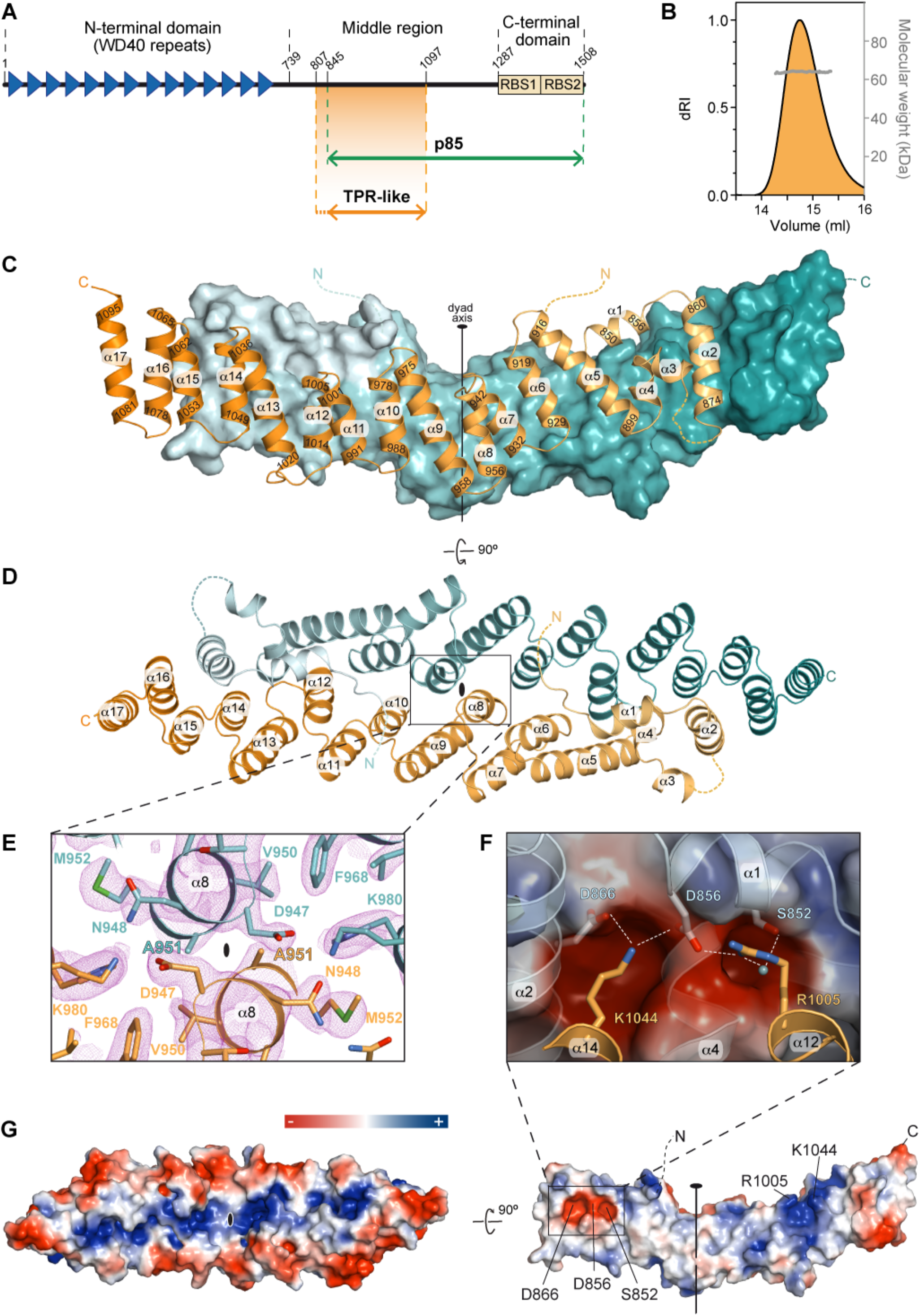
Crystal structure of Gemin5 TPR-like dimerization module. **(A)** Scheme of Gemin5 protein. WD40 repeats and RBS domains are indicated with blue triangles and yellow boxes, respectively. The p85 fragment, resulting from L protease cleavage during viral infection, and the TPR-like domain are indicated with green and orange double arrows, respectively. **(B)** SEC-MALS analysis proves that G5-TPR is a dimer in solution, with a molecular weight of 64 kDa (± 0.01%). **(C)** Crystal structure of the G5-TPR dimer with the subunit in front represented in cartoon and the subunit at the back shown in surface representation. The subunits are depicted in different color from lighter to darker from the N-to the C-terminus. Dashed lines indicate regions not seen in the electron density maps. Numbers indicate the residues at the N- and C-ends of α-helices. **(D)** Perpendicular (“top”) view of **(C)** with both subunits represented in cartoon. The position of the dyad axis is indicated in **(C)** and **(D)**. **(E)** Detail of the intersubunit interactions across the dyad axis and localization of residue A951. The 2F_obs_-F_calc_ electron density map at 1 σ is shown in pink mesh. **(F)** Electrostatic interactions between residues D866 and D856 in one subunit (shown in semi-transparent surface representation) and R1005 and K1044 in the other subunit (shown as orange cartoon). A water molecule bridging the side chains of R1005 and S852 is shown as a cyan sphere. **(G)** Surface representation of the protein dimer (left) and of the dimerization interface of one of the subunits (right), colored according to the electrostatic potential. Blue and red contours begin at +2.5 and −2.5 kT, respectively. See also Figures S1–S5.

Gemin5 contains two distinct functional regions at the protein ends (Figure 1A). The N-terminal part (amino acids 1–739) is composed of two juxtaposed seven-bladed WD40 domains (Jin et al., 2016) that recognize the Sm-site of snRNAs and the m^7^G cap via base-specific interactions (Tang et al., 2016; Xu et al., 2016). The C-terminal part (amino acids 1287–1508) on the other hand, harbors a non-canonical RNA-binding site (RBS), with two moieties designated as RBS1 and RBS2 (Fernandez-Chamorro et al., 2014), which differ in RNA-binding capacity as well as in the ability to modulate selective translation. NMR structural analysis of the RBS1 polypeptide revealed a mixture of conformations in solution (Fernandez-Chamorro et al., 2014), frequently found in unstructured protein domains (Jonas and Izaurralde, 2013). However, the biological relevance of the ∼550 amino acids bridging the two functional ends of Gemin5 remains elusive.

Here, to better understand the potential role of the middle region of Gemin5, we isolated a fragment of the protein spanning amino acids 807–1097 and demonstrated that it forms a stable homodimer in solution. The crystal structure revealed a compact elongated canoe-shaped dimer with each subunit folding into a tetratricopeptide repeat (TPR)-like domain. Based on the structure, we designed a mutation A951E that disrupts the formation of the dimer. We further proved by pull-down and mass spectrometry analysis that the cleavage product p85 wild-type (wt) but not the p85 bearing mutation A951E interacts with Gemin5 in living cells. These data also revealed that p85-wt, but not p85-A951E, recruits cellular proteins involved in RNA splicing and translation processes. Furthermore, while p85-wt stimulated IRES- and cap-dependent translation, p85-A951E failed to enhance translation in human cells. Together, our results uncover a crosstalk between the ability of the dimerization module to associate with the full-length protein and the capacity of p85 to modulate translation.

## RESULTS

### The middle region of Gemin5 bears a novel dimerization domain

A comprehensive bioinformatic analysis of the middle region of Gemin5 predicted the existence of an α-helix rich domain comprising residues 807–1097 with highest similitude to the tetratricopeptide repeat (TPR)-like domain of the elongator complex protein 1 (Elp1) (Xu et al., 2015). Guided by the predictions, we produced in bacteria a construct covering the putative TPR-like region of human Gemin5 (residues 807–1097) (Figure 1A) fused at the N-terminus to a cleavable polyhistidine-tagged maltose binding protein (MBP). The protein without tag and purified to homogeneity was highly soluble and the analysis by size-exclusion chromatography coupled to multi-angle light scattering (SEC-MALS) indicated a molecular mass of 64 kDa (Figure 1B). This is twice the expected 32 kDa mass for a single copy of the protein, indicating that Gemin5 residues 807–1097 form a stable homodimer in solution.

Diffraction quality crystals grown in presence of sodium iodide allowed to solve the structure of the protein at 2.7 Å resolution using single-wavelength anomalous dispersion (SAD) (Table 1). The protein model was used for phasing by molecular replacement a second data set collected up to 2 Å resolution from an isomorphous crystal grown with tartrate instead of iodide (Table 1). The crystal structure revealed a dimer formed by two molecules of the protein related by a twofold non-crystallographic symmetry (Figures 1C–1E). The two subunits in the dimer are similar (Figure S1), and consist of 17 α-helices forming a flat and elongated right-handed α-solenoid with a 40° angle kink between helices α1–8 on one side and helices α9–17 on the other (Figures 1C and 1D). This helical palisade is nucleated by six antiparallel double-helix repeats (helices α5-6, α7-8, α9-10, α11-12, α13-14 and α15-16) with structural and sequence similarity to the classical PP5 TPR motif (Das et al., 1998) (Figure S2). A C-terminal “capping” helix (α17) (Figures 1C and 1D), present in most TPR domains (D’Andrea and Regan, 2003), covers the hydrophobic surface of the last helical repeat at one end of the subunit. At the other end, the structure is completed by two additional N-terminal helical pairs: helix α1 lays perpendicular on “top” of the palisade and makes a 90° angle with helix α2, which connects through a disordered region (residues 877–883) to a pair of short α-helices, α3 and α4, also related by a 90° angle (Figures 1C and 1D). There was no electron density attributable to the N-terminal residues 807–844, and thus, these 38 amino acids were not included in the final models. Indeed, we noticed that the protein reproducibly experimented a proteolytic degradation, yielding a ∼28 kDa product (Figure S3) that agrees in size with the crystal model, suggesting that the N-terminal sequence is cleaved off during crystallization. Interestingly, the cleavage site coincides with the ^843^RKAR^846^ motif used by the L protease to release the p85 polypeptide during foot-and-mouth disease virus (FMDV) infection (Figure 1A) (Pineiro et al., 2012).

**Table 1.** Data collection and refinement statistics.

|  | **G5-TPR (SAD)** | **G5-TPR (native)** |
| --- | --- | --- |
| PDB ID | 6RNS | 6RNQ |
| **Data Collection** |  |  |
| Beamline | BL13-XALOC, ALBA | BL13-XALOC, ALBA |
| Wavelength (Å) | 2.0330 | 0.9792 |
| Space group | P2<sub>1</sub> | P2<sub>1</sub> |
| Cell dimensions<br>*a*, *b*, *c* (Å)<br>$\alpha$, $\beta$, $\gamma$ (°) | 67.8, 54.6, 103.0<br>90, 108.8, 90 | 64.2, 55.0, 103.6<br>90, 104.3, 90 |
| Total reflections | 63,764 (9,179)<sup>a</sup> | 172,756 (25,023) |
| Unique reflections | 19,865 (2,881) | 51,344 (7,460) |
| Resolution (Å) | 64.22 - 2.69 (2.84 - 2.69) | 100.43 - 1.95 (2.05 - 1.95) |
| R<sub>meas</sub> | 0.162 (1.475) | 0.074 (1.461) |
| Mean I/σI | 6.4 (1.0) | 9.1 (1.0) |
| Completeness (%) | 98.8 (98.9) | 99.6 (99.6) |
| Redundancy | 3.2 (3.2) | 3.4 (3.4) |
| CC<sub>1/2</sub> | 0.99 (0.75) | 0.99 (0.67) |
| **Refinement** |  |  |
| Resolution (Å) | 64.22 - 2.69 | 100.4 - 1.95 |
| No. of reflections | 19,722 (1,939) | 51,194 (5,068) |
| R<sub>work</sub>/R<sub>free</sub> | 0.23 / 0.26 | 0.21 / 0.23 |
| Rmsd<br>Bond lengths (Å)<br>Bond angles (°) | 0.004<br>0.84 | 0.011<br>1.13 |
| No. of atoms<br>Protein<br>Ligand<br>Water | 3,805<br>13<br>26 | 3,819<br>1<br>74 |
| Ramachandran plot<br>Favored (%)<br>Allowed (%)<br>Outliers (%) | 96.03<br>3.97<br>0.00 | 97.93<br>2.07<br>0.00 |

<sup>a</sup> Values in parenthesis correspond to the highest resolution shell.

The two protein subunits are confronted in antiparallel orientation, forming an elongated canoe-shaped homodimer approximately 120 Å long x 35 Å wide, with helices α16 and α17 from one subunit overtaking the other subunit at the stern and bow ends (Figures 1C and 1D). The dimer is formed by the intertwined α-helices in the second position of the helical repeats, with contributions of side chains from the helices in the first position of the repeats, and also by the N-terminal helices α1, α2 and α4 (Figure 1D). The closest intersubunit distance occurs between the two A951 residues at helices α8 flanking the molecular dyad axis (Figures 1D and 1E). Overall, the dimerization interface buries ∼3,400 Å^2^ per subunit as computed by PISA (Krissinel and Henrick, 2007), which accounts for 24% of the total protein surface and suggests the formation of a stable dimer. This tight association occurs mostly through hydrophobic residues but also includes thirteen hydrogen bonds and two pairs of salt bridges between patches of complementary charge formed by residues R1005 and K1044 (both in α12) in one subunit and D856 (α1) and D866 (α2) in the other (Figure 1F). The solvent exposed area is hydrophilic, with a positively charged narrow groove on “top” of the dimer surface delimited by a rim of acidic residues (Figure 1G).

A search with the DALI server (Holm and Sander, 1995) confirmed the structural similarity between the TPR-like domain of Gemin5 (hereafter named as G5-TPR) and other TPR-containing proteins, particularly with Elp1 (Xu et al., 2015), showing highest Z-score of 11.1 and RMSD of 5.2 Å for 200 Cα positions (Figures S4A and S4B).

### Mutation of a conserved residue at the dyad axis abrogates protein dimerization

The alignment of 145 Gemin5 sequences from vertebrates shows an identity in the TPR-like region of 55%, which increases to 86% within mammals (Figure 2A). The most conserved residues are involved in interactions within the subunit and across the dimer interface (Figures 2B and 2C). These results strongly suggest that the dimerization module is an evolutionary preserved feature of Gemin5.

**Figure 2.**
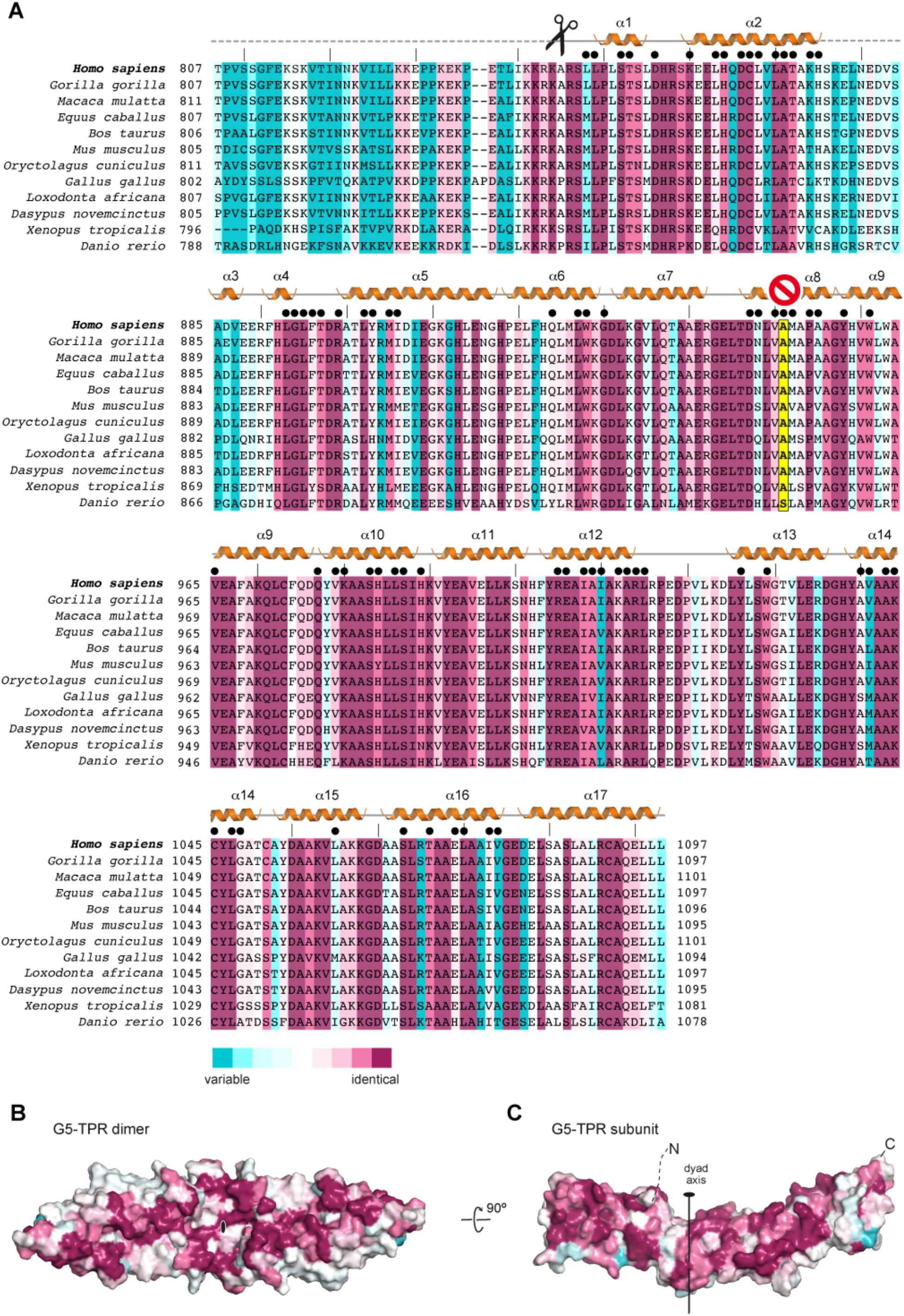
Conservation of the dimerization module in Gemin5. **(A)** Alignment of 145 Gemin5 sequences spanning the TPR-like domain, out of which only twelve are represented for simplicity. Residues are colored according to their conservation, from magenta (identity) to cyan (variable). The α-helices of human G5-TPR are depicted above the sequence, and missing regions in the model are indicated with dashed lines. The scissors mark the L protease cleavage motif. Residue A951, whose substitution by glutamate impedes human G5-TPR dimerization, is indicated with a yellow background and a prohibition sign, whereas residues involved in intersubunit interactions are denoted with a black dot on top. **(B,C)** Sequence conservation plotted on the surface of the protein dimer **(B)** and on the dimerization interface **(C)**. The molecules are represented in the same orientation as in Figure 1G.

To further study the functionality of this association, we attempted to disrupt the dimerization of the G5-TPR introducing a glutamate at the central position occupied by residue A951 (Figures 1E and 2A). The larger and negatively charged side chain of glutamate was expected to repel the two subunits due to steric clashes and charge repulsion. Although the G5-TPR A951E variant was successfully expressed in bacteria, removal of the N-terminal MBP tag caused the protein to precipitate heavily under conditions in which the wild-type G5-TPR is soluble up to 15 mg ml^−1^ (Figure S5). These results indicate that mutation A951E disrupts the dimerization of G5-TPR, and that the isolated subunit is not stable in solution.

### The Gemin5 dimerization domain lacks RNA-binding ability

Since the functions reported for Gemin5 are directly connected to RNA-dependent pathways (Francisco-Velilla et al., 2016; Yong et al., 2010), we sought to investigate whether or not the dimerization module interacts with RNA. The RNA binding capacity of the purified G5-TPR was assessed in parallel to the RBS1 domain of Gemin5, previously shown to interact with domain 5 (d5) of the FMDV IRES (Fernandez-Chamorro et al., 2014), as well as to recognize multiple RNA targets in human cells (Francisco-Velilla et al., 2018). As shown in Figure 3A, G5-TPR was unable to form a retarded complex with d5 RNA, even at high protein concentration (1 µM), while RBS1 exhibited a robust RNA-binding affinity. To reinforce this result, we analyzed the capacity of the G5-TPR to interact with RNAs differing in sequence and secondary structure. All these RNAs (Figures 3B–3D) [a hairpin of 22 nt, a 180 nt RNA adopting a complex stem-loop structure (Francisco-Velilla et al., 2018) and a single stranded RNA of 32 nt (Pineiro et al., 2013)] failed to form a retarded complex with the purified protein at high concentrations (1 µM). Furthermore, gel-shift assays carried out with ssDNA (29 nt) or dsDNA (29 bp) indicated that G5-TPR (up to 4.3 µM) does not recognize DNA (Figures 3D and 3E). Therefore, we conclude that the Gemin5 dimerization domain does not promote interaction with nucleic acids in vitro.

**Figure 3.**
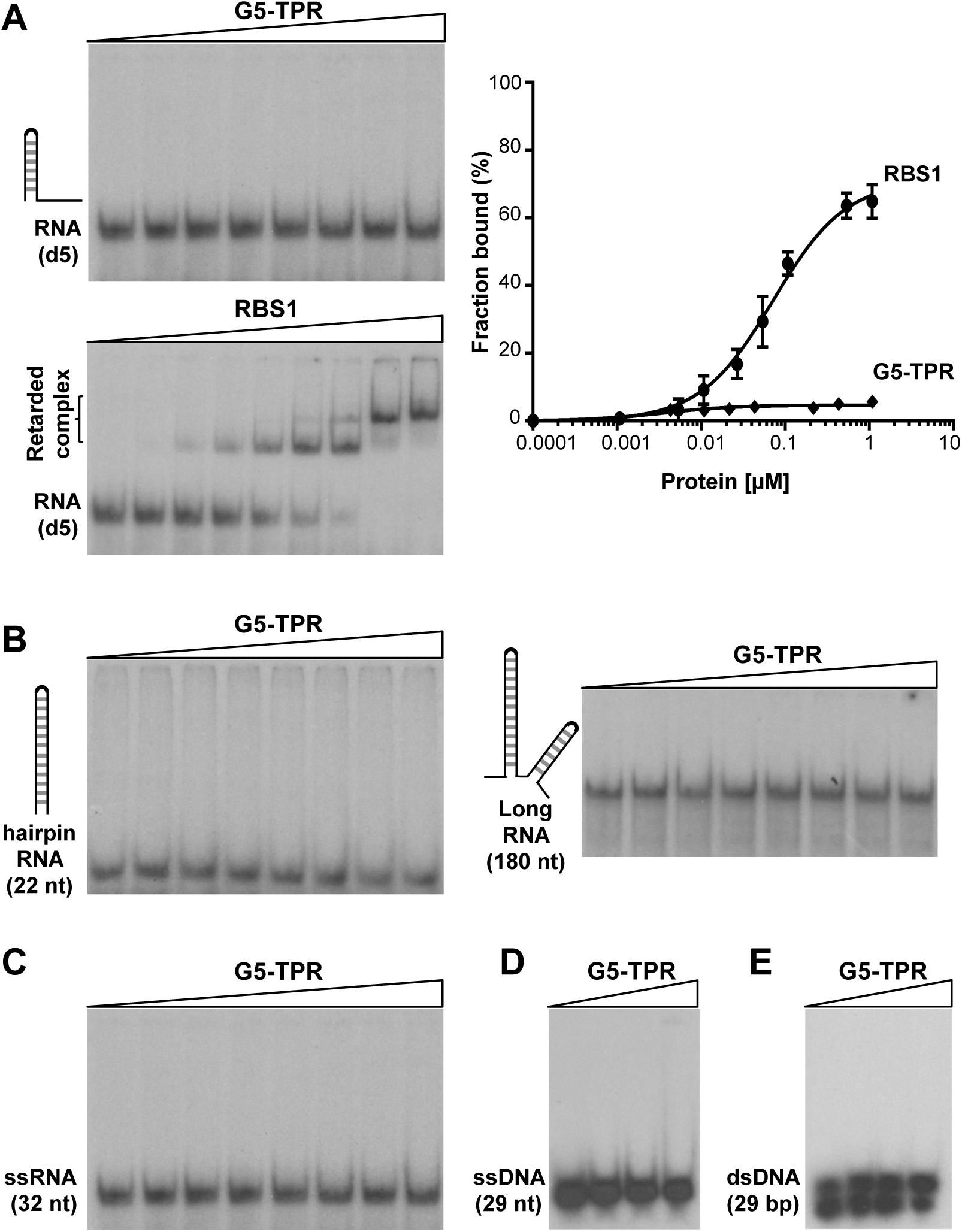
G5-TPR lacks nucleic acids binding capacity. **(A)** Representative example of a gel-shift assay conducted with increasing amounts of purified G5-TPR (0 to 1 µM) and labeled domain 5 (d5) RNA. Purified RBS1 (0 to 2.4 µM) was used as a positive control of the RNA-binding assay. The percentage of retarded complexes obtained in three independent assays is plotted on the right panel. Values represent the mean ± SEM. **(B,C)** RNA-binding assays performed with G5-TPR and various probes: a 22 nt hairpin RNA, a 180 nt RNA predicted to fold in several stem-loops and a single stranded 32 nt RNA. **(D,E)** Lack of retarded complex formation with ssDNA (29 nt) or ds DNA (29 bp).

### The Gemin5 cleavage product p85, but not p85-A951E, recruits the full-length protein in living cells

The finding of a novel dimerization module in the middle region of Gemin5 prompted us to study the functional relevance of this protein-protein interaction. Previous data showed the immunodetection of the endogenous Gemin5 in TAP-pull-down protein complexes isolated with a Gemin5 fragment spanning residues 1–1287 (Francisco-Velilla et al., 2016), suggesting that Gemin5 could oligomerize in the cell cytoplasm. However, the protein region responsible for this association was unknown. On the other hand, we have shown that Gemin5 is proteolytically cleaved in infected cells generating the polypeptide p85 (Pineiro et al., 2012), which includes the dimerization module (Figure 1A). Hence, we posit that the physiological relevance of the dimerization module could be linked to p85 function.

To determine whether p85 associates with full-length Gemin5 through the dimerization module in a cellular environment, we performed mass spectrometry analysis of proteins copurifying with p85-wt-TAP expressed in HEK293 cells (Dataset 1). Gemin5 was unequivocally identified in two replicates [score 375.86 and 344.37, coverage 55% and 42%, respectively]. Besides numerous peptides from the p85 region, analysis of Gemin5 amino acid sequences revealed the presence of trypsin fragments corresponding to residues 1–844, yielding a total of 33 peptides (Figures 4A and S6). Hence, these results show that the full-length Gemin5 copurifies with p85-wt-TAP, reflecting the formation of a stable complex with the endogenous protein.

**Figure 4.**
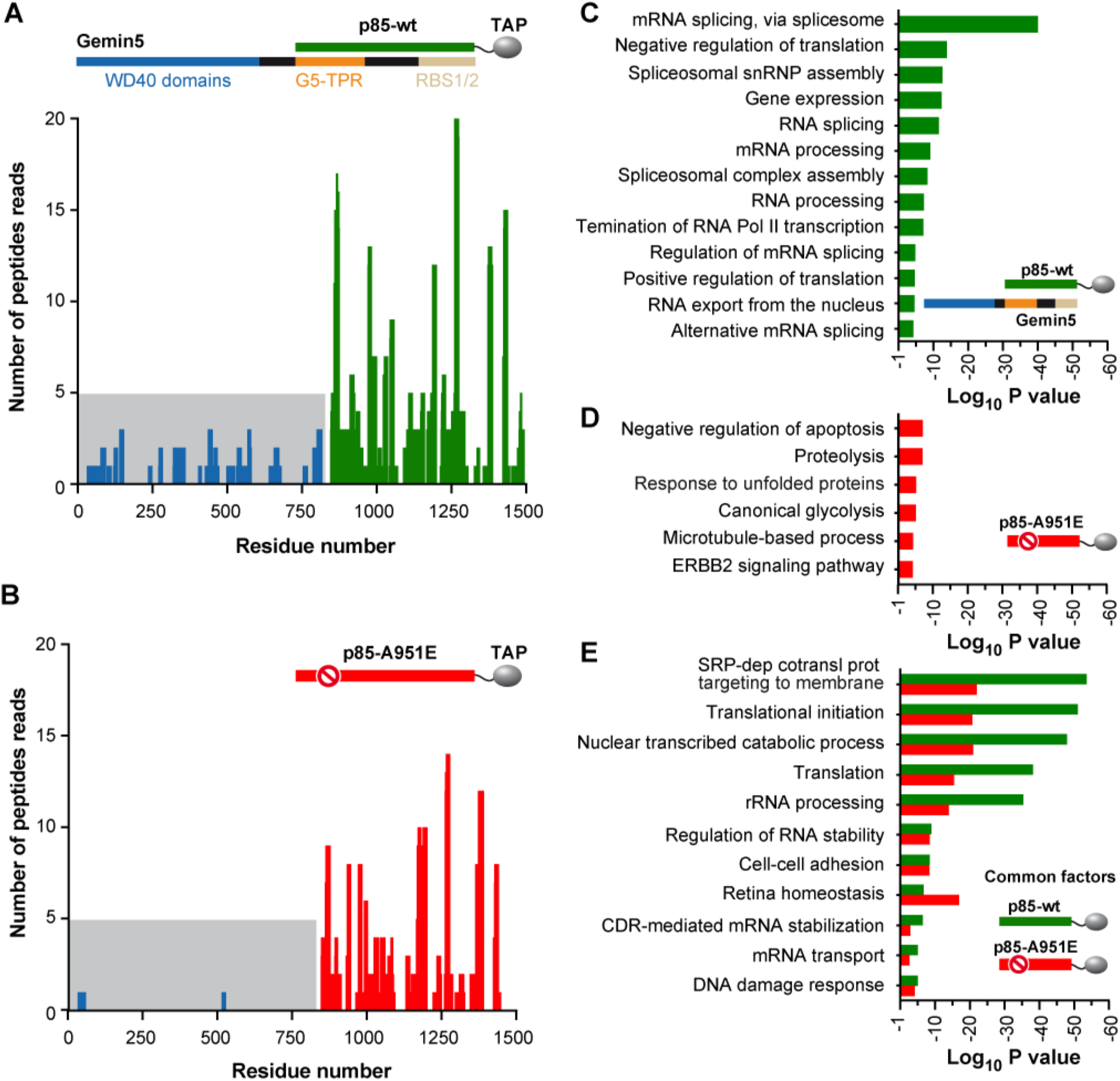
The p85 region of Gemin5 retrieves the endogenous Gemin5 in living cells. HEK293 cells were transfected with plasmids expressing p85-wt-TAP **(A)** or the mutant p85-A951E-TAP **(B)**. Twenty-four hours later, cells were lysed, and the proteins copurifying with p85-wt or p85-A951E were identified by mass spectrometry. The number of peptide reads corresponding to Gemin5 determined in TAP samples of p85-wt **(A**; green bars) and p85-A951E **(B**; red bars) are represented according to their position. Peptide reads corresponding to Gemin5 residues 1-844 are colored in blue and highlighted in grey background. **(C,D)** Gene ontology (GO) classification obtained for the overlap of the pulldown replicates obtained with p85-wt only (**C**; green), or p85-A951E only (**D**; red). The top GO terms are represented according to P value; cut-off was set to 10^−5^. The legend depicts p85-Gemin5 association; p85-A951E fails to capture Gemin5. **(D)** GO terms shared by proteins p85-wt and p85-A951E. See also Figures S6 and S7.

To further investigate the biological relevance of the Gemin5 dimerization module we carried out mass spectrometry analysis of cellular proteins associated with p85-A951E-TAP mutant (Dataset 1). Analysis of the peptides copurifying with p85-A951E-TAP (score 330.38 and 292.0 for the biological replicate samples, similar to p85-wt-TAP) revealed only 4 peptides within the region 1–844 of Gemin5 (Figures 4B and S7), in contrast to the 33 peptides obtained for p85-wt-TAP. Hence, we conclude that the G5-TPR domain is necessary for the dimerization of Gemin5, and that mutation A951E disrupts the capacity of p85 to recruit the full-length protein in living cells.

### The functional groups retrieved by p85 uncover the physiological relevance of Gemin5

Next, we sought to investigate the implication of protein-protein interactions on the functional groups copurifying with either p85-wt, which recruits the endogenous Gemin5 (Figure 4A), or p85-A951E, unable to do so (Figure 4B). The Gene Ontology (GO) annotation of the biological processes obtained with DAVID database (Huang da et al., 2009) for the overlap of the biological replicates was classified according to P value for three separate groups: i), proteins copurifying with p85-wt, which retrieves the full-length Gemin5 (Figure 4C); ii), proteins exclusively copurifying with p85-A951E (Figure 4D); and iii), proteins shared by p85-wt and p85-A951E, presumably interacting with residues 845–1508 (Figure 4E). The top GO functional groups copurifying exclusively with p85-wt are involved in mRNA splicing (10^−40^), negative regulation of translation (10^−14^), and snRNP assembly (10^−13^) (Figure 4C). These results are fully consistent with the functional properties reported for Gemin5 (Francisco-Velilla et al., 2019). This is followed by 10 additional GO groups related to RNA splicing, processing and translation (gene expression, regulation of RNA splicing, mRNA processing, positive regulation of translation, and RNA export; P values ranging between 10^−13^ to 10^−5^).

In marked difference with p85-wt, none of the GO terms related to splicing and translation members, were detected with the mutated p85-A951E protein (Figure 4D). In addition, the number of GO groups exclusively copurifying with the A951E mutant were fewer, and had lower statistical significance, although the protein was identified with similar score. Remarkably, p85-A951E copurified with members of the protein degradation pathway since the top GO terms are involved in negative regulation of apoptotic processes (10^−7^), proteolysis (10^−7^), and response to unfolded protein (10^−6^), suggesting protein misfolding. None of these terms were observed with the p85-wt protein.

Besides the GO groups specific for the wt or the mutant version of p85, a large number of factors copurified with both proteins (Figure 4E). Thereby, these factors mostly interact with residues 845–1508 of Gemin5. GO terms shared by both p85-wt and p85-A951E do not involve RNA splicing, revealing distinct functional properties of the Gemin5 domain expressed in cells. According to P value, the top GO term was SRP-dependent cotranslational protein targeting to membrane (P values for p85-wt and A951E 10^−54^ and 10^−22^, respectively), suggesting that Gemin5 interacts through the 845–1508 region with SRP (signal recognition particle). This result is in full agreement with a report showing that the Gemin5 alone was capable of interacting with SRP particle (Piazzon et al., 2013). The next interacting groups are involved in translation initiation (P value 10^−52^ wt, 10^−21^ A951E), nuclear transcribed metabolic processes, translation, and rRNA processing (P values ranging between 10^−48^ to 10^−36^ and 10^−21^ to 10^−14^ for wt and A951E, respectively), consistent with recently reported features of the protein (Francisco-Velilla et al., 2018; Francisco-Velilla et al., 2016; Garcia-Moreno et al., 2019). To our surprise, GO terms with significant P values are involved in regulation of RNA stability, cell-cell adhesion, retina homeostasis, coding region instability determinant (CRD)-mediated mRNA stabilization, mRNA transport, and DNA damage response, suggesting the involvement of Gemin5 in still unknown functions. In summary, there is a crosstalk between the ability of p85 to recruit Gemin5, driven by the dimerization module, and its capacity to associate with partners directly involved in RNA splicing, RNA processing, and translation control.

### Disruption of the dimerization ability correlates with loss of translation stimulation by p85

As we and others have shown, Gemin5 is involved in translation control (Francisco-Velilla et al., 2018; Pacheco et al., 2009; Workman et al., 2015). However, while the full-length protein down-regulates translation, the p85 product observed in infected cells enhances IRES-dependent translation (Fernandez-Chamorro et al., 2014). Functional assays carried out in HEK293 cells co-expressing Xpress-tagged p85 and luciferase downstream of the FMDV IRES (Figure 5A) revealed that expression of Xpress-p85-wt concurred with IRES-dependent stimulation of luciferase activity (Figure 5B) (P = 0.027). In contrast, similar levels of expression of the Xpress-p85-A951E construct failed to do so (Figure 5B) (P = 0.404). Likewise, the stimulation of cap-dependent translation observed in p85-wt was not detected in the mutant p85-A951E (Figure 5B) (P = 0.013 and 0.157, respectively). Moreover, to determine whether the region comprising RBS1/2 was essential for translation control, we used a p85 construct deleting the C-terminal region up to amino acid 1287 (Figure 5A). Expression of Xpress-p85ΔRBS1/2 resulted in lack of effect on IRES- and cap-dependent translation regulation (Figure 5C) (P = 0.180 and 0.856, respectively). Thus, we conclude that the dimerization domain is necessary but not sufficient to enhance translation. Therefore, the ability to modulate translation requires the presence of the entire p85 region, including the RBS domains at the C-terminus.

**Figure 5.**
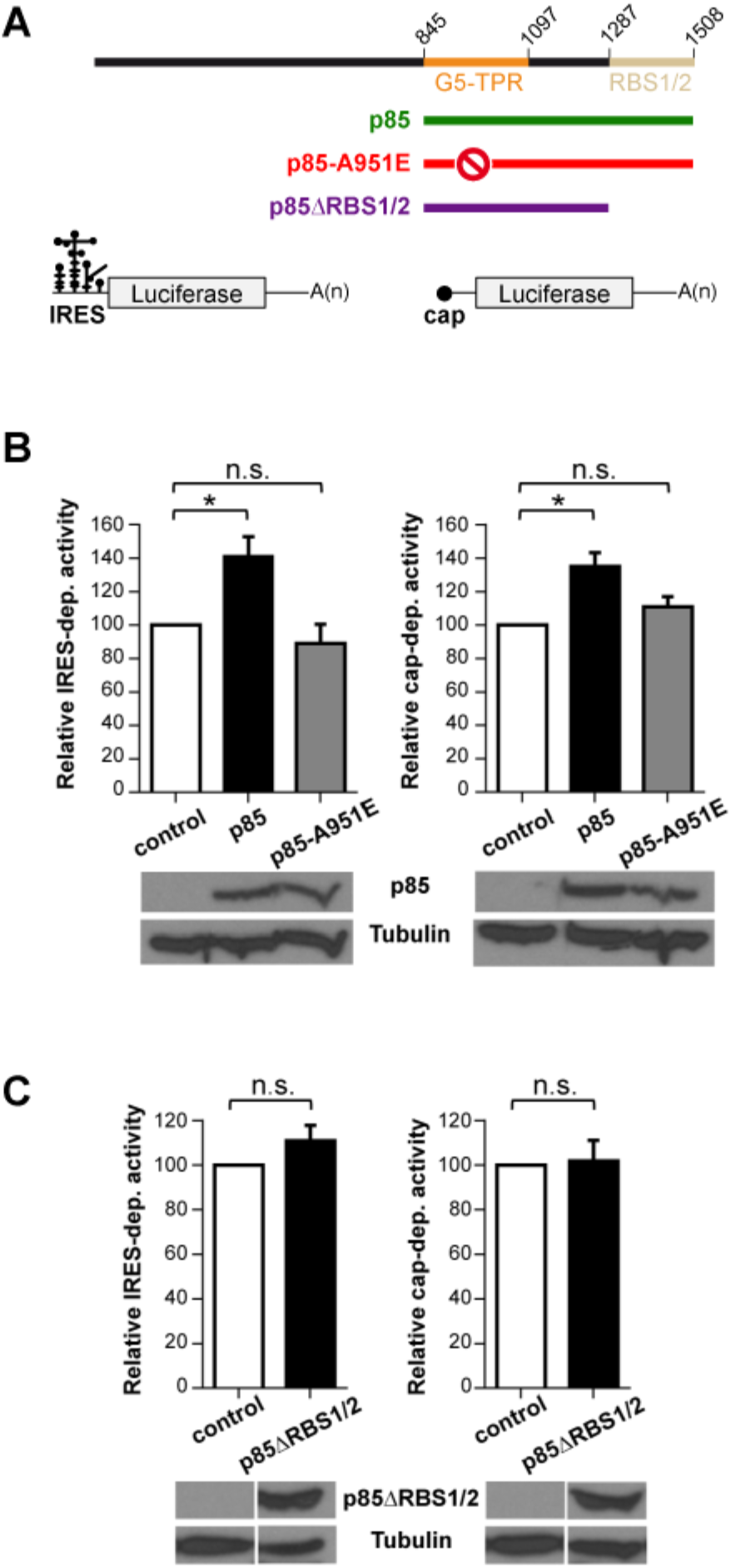
Translation stimulation by p85 depends on the TPR dimerization domain and the RNA-binding domain. **(A)** Diagram of the p85, p85-A951E and p85ΔRBS1/2 proteins expressed with the Xpress-tag (top), and of the luciferase reporter mRNAs with IRES or cap sequences (bottom). **(B)** Luciferase activity measured in HEK293 cell lysates expressing IRES-mRNA or cap-mRNA co-transfected with Xpress-p85 and Xpress-p85-A951E. Expression of Xpress-p85 and Xpress-p85-A951E was monitored by western blot using anti-Xpress. Tubulin was used as loading control**. (C)** Luciferase activity assay using Xpress-p85ΔRBS1/2 construct. Protein expression was monitored by western blot using anti-Xpress. In all cases, luciferase values are normalized to cells expressing the empty vector conducted side by side. Values represent the mean ± SEM obtained in three independent assays. Asterisks denote *p*-values (* P < 0.05).

## DISCUSSION

In this study, we combined structural, proteomic and functional analysis to get insights into the multitasking protein Gemin5. We demonstrate that the middle region of Gemin5 contains a previously uncharacterized dimerization module, which also remains at the N-terminus of the viral-induced cleavage fragment p85 (Figure 6A). The TPR-like motifs of Gemin5 do not conform to the archetypical length of 34 amino acids, but rather alternate helical pairs of 32-33 and 27-29 amino acids that exhibit the loosely conserved TPR pattern of small and large hydrophobic residues (D’Andrea and Regan, 2003; Zhu et al., 2016) (Figure S2). However, whereas most tandem TPR arrays assemble into a right-handed super helical structure with concave and convex surfaces for protein-protein and protein-RNA interactions (Kajander et al., 2007), the helical repeats in Gemin5 are stretched in an α-solenoid rod that provides an extensive flat surface for self-dimerization. Importantly, and despite the tight association between G5-TPR subunits, a single point mutation in the dyad axis, A951E, was sufficient to disrupt the dimer (Figures 1E and S3).

**Figure 6.**
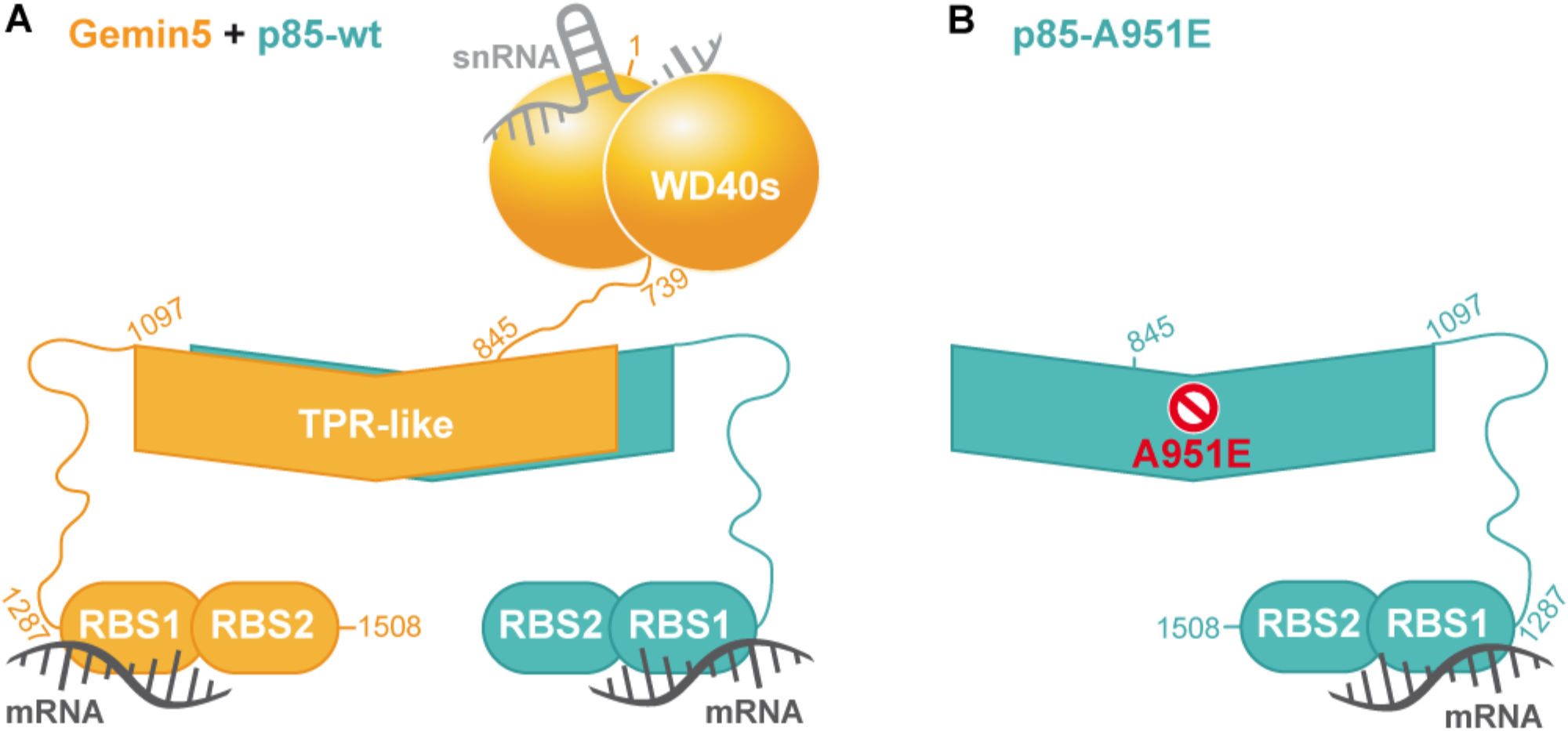
Model of Gemin5-p85 heterodimer driven by the TPR-like domain. **(A)** Schematic representation of the association between Gemin5 (orange) and p85 (greenish). The relative orientation between the WD40s, TPRs and RBS1/2 domains of Gemin5 is unknown, although they are represented at three different levels to highlight their distinct roles in snRNA and ribosome binding (WD40s), dimerization (TPR-like) and RNA translation (RBS1/2). **(B)** Schematic representation of p85-A951E. The prohibition sign indicates that mutation A951E impedes the interaction with Gemin5.

The canoe-shaped conformation of the G5-TPR homodimer is singular, and an exhaustive manual and computational search failed to retrieve other TPR motifs resulting in a similar association. Although a search with DALI highlighted the structural similarity between G5-TPR and the C-terminal region of Elp1, most interactions in the Elp1 homodimer occur between the TPR-like motifs in one subunit and the “capping” helical bundle in the other (Xu et al., 2015) (Figure S4B). Instead, 14 out of the 17 α-helices contribute directly to G5-TPR dimerization (Figures 1C, 1D, S2, and S4A). The dimerization mode of Gemin5 is more reminiscent of the association between the yeast vesicle coating proteins β’-COP and α-COP of the COPI complex (Lee and Goldberg, 2010), and Sec31 of the COPII complex (Fath et al., 2007). These proteins present α-solenoid regions that in α-COP and β’-COP interact in an antiparallel orientation with a 40° angle, forming a stable heterodimer (Figure S4F) while in Sec31, the central α-solenoid regions of two subunits interact about a 2-fold axis, creating an interlocked homodimer (Figure S4G). Curiously, alike in Gemin5, the α-solenoids of the mentioned proteins are preceded by N-terminal WD40 domains (Figures 1A and 6A), highlighting how a similar arrangement of these domains evolved into proteins with different functions.

We also show here that the G5-TPR alone does not interact with RNA, irrespectively of whether it consisted of single or double stranded molecules, short hairpins, long stem-loops, or DNA (Figure 3). This result establishes a main difference with proteins carrying TPR domains involved in RNA-dependent pathways or in direct RNA-binding (Abbas et al., 2013; Grotwinkel et al., 2014; Johnson et al., 2018; Katibah et al., 2013; Yang et al., 2012; Zhou et al., 2018). The high sequence conservation of the TPR-like domain strongly suggests that the dimerization module of Gemin5 plays a fundamental role for the architecture and activity of the protein. Interestingly, the first residue detectable on the crystal structure occurs immediately after residue K844, matching the L protease cleavage site that releases the p85 fragment during viral infection (Pineiro et al., 2012). Consequently, the dimerization module might play a central structural role also in the p85 product. Indeed, we have found that in living cells, full-length Gemin5 is captured with a p85 construct with a TAP-tag at the C-terminus (Figures 4A, 6A and S6). However, a p85-TAP variant carrying the point mutation A951E that prevents dimerization failed to recruit the endogenous Gemin5 protein (Figures 4B, 6B and S7). These data demonstrate that the TPR-like region is necessary for the interaction of p85 with the available Gemin5 protein in the cell environment, presumably contributing to the behavior of this viral-induced cleavage fragment.

Here we provide direct evidence for physical p85-Gemin5 interactions (Figure 4A), which in turn, unveiled fundamental properties of Gemin5. The top GO terms retrieved by p85-wt, which establishes protein-protein bridges with the endogenous Gemin5, are splicing-related components, translation control members, and snRNP assembly factors (Figure 4C). All these groups are consistent with the functions reported for this protein (Francisco-Velilla et al., 2019), and suggest that these components probably interact with the N-terminal region of Gemin5. In contrast, GO functional groups shared by p85-wt and p85-A951E constructs (thereby, factors shared with the C-terminal moiety 845–1508) are related to SRP-dependent cotranslational protein targeting to membrane, translation initiation and nuclear transcribed catabolic processes (Figure 4D). These families of factors confirm the known roles of Gemin5 and strongly suggest new potential functions for this versatile protein. Finally, GO terms associated uniquely to p85-A951E, which fails to retrieve the endogenous Gemin5 protein, are related to apoptotic processes, proteolysis, and response to unfolded proteins. We attribute this result to a misfolding problem of the protein caused by the destabilizing mutation. Together, these results corroborate that different domains of Gemin5 are responsible for its multiple functions, and highlight the biological relevance of the dimerization module of this protein.

The properties unveiled in our study of the dimerization module of Gemin5 shed new light on this little characterized protein. We show here that the TPR-like polypeptide alone does not recognize RNA in vitro (Figure 3), and also, that this region of the protein is not sufficient to regulate translation in living cells (Figure 5C). However, disruption of the complex formation by mutation A951E abolishes the translation enhancing effect of p85 (Figure 5B). These results indicate that p85 comprises additional determinants for RNA-dependent pathways. Importantly, p85 harbors the bipartite non-canonical RBS1/2 domain at its C-terminus (Fernandez-Chamorro et al., 2014). The RBS1 domain not only displays robust RNA-binding capacity in vitro (Figure 3A), but more importantly this region of Gemin5 binds selective RNA targets in living cells (Francisco-Velilla et al., 2018). Therefore, the translation stimulatory capacity of p85 reflects the combined properties of the dimerization module and the RBS domains (Figures 6A and 6B).

Overall, these results led us to propose that mutations in the dimerization module of Gemin5 will result in loss of function. The protein is broadly expressed in all human tissues (Kim et al., 2014; Uhlen et al., 2015), and the loss of Gemin5/*rigor mortis* in *Drosophila* is lethal (Gates et al., 2004). On the other hand, altered levels of SMN proteins causing defects in SMN complex assembly lead to spinal muscular atrophy (SMA), a severe child disease (Burghes and Beattie, 2009). According to our data, mutation A951E disrupts the p85-Gemin5 interaction, hampers the association with proteins regulating splicing and translation processes, and abolishes translation control. Thus, it is plausible that defects in Gemin5 dimerization will course with severe disease. In support of this possibility, mutations affecting the TPR domains of other proteins have been shown to be functionally critical. Thus, splicing variants producing a protein lacking the TPR-like dimerization domain of Elp1 are involved in > 95% of all familial dysautonomia patients (Xu et al., 2015), whereas nonsense mutations in the TPR region of aryl-hydrocarbon-interacting protein-like 1 (AIPL1) cause inherited retinopathies (Sacristan-Reviriego et al., 2017), and missense mutations in the TPR domain of O-linked N-acetyl-glucosamine transferase cause intellectual disability (Gundogdu et al., 2018).

In summary, we have disclosed an unprecedented dimerization module in Gemin5 and provided direct evidence for physical p85-Gemin5 interactions in the cell and its relationship to translation control. Further work will be needed to explore the implications of Gemin5 dimerization in other activities of this multifaceted protein.

## ACKNOWLEDGEMENTS

This work was supported by MINECO (grants BFU2016-80570-R, BFU2017-84492-R, AEI/FEDER, UE), Comunidad de Madrid (B2017/BMD-3770) and an Institutional grant from Fundación Ramón Areces. X-ray diffraction experiments were performed at XALOC beamline at ALBA Synchrotron with the collaboration of ALBA staff, with special thanks to Fernando Gil. We thank the CBMSO Proteomics Unit for help with the MS/MS analysis, the CNIO Spectroscopy Unit for SEC-MALS analysis, Jorge Ramajo for technical assistance, and Crisanto Gutierrez for valuable comments on the manuscript.

## AUTHOR CONTRIBUTIONS

MM-M, SR-M, RF-V, EM-S designed the experiments, MM-M, JF-C performed the protein purification, MM-M, SR-M collected X-ray diffraction data and determined the protein structure, MM-M performed SEC-MALS measurements, AE-B performed the EMSA binding assays, RF-V, AE-BV performed the protein-TAP experiments, and analyzed the proteomic data, RF-V performed the p85-dependent translation assays in human cells, MM-M, SR-M, EM-S wrote the paper with comments from all authors.

## STAR METHODS

PROTEIN ENGINEERING FOR STRUCTURAL AND FUNCTIONAL STUDIES

EXPRESSION AND PURIFICATION OF G5-TPR

GEL-FILTRATION COUPLED TO MULTI-ANGLE LIGHT-SCATTERING (SEC-MALS) MEASUREMENTS

CRYSTALLIZATION

DATA COLLECTION AND STRUCTURE DETERMINATION

RNA, DNA ELECTROPHORETIC MOBILITY SHIFT ASSAY

PROTEIN COMPLEXES PURIFICATION BY TANDEM AFFINITY PURIFICATION (TAP)

IN-GEL DIGESTION and MASS SPECTROMETRY IDENTIFICATION

GENE ONTOLOGY ANALYSIS

GEMIN5 POLYPEPTIDES EXPRESSION AND LUCIFERASE ACTIVITY ASSAYS

IMMUNODETECTION

STATISTICAL ANALYSES

## Methods

### Protein engineering for structural and functional studies

The bioinformatic servers HHPred (Zimmermann et al., 2018), Phyre2 (Kelley et al., 2015) and I-TASSER (Yang et al., 2015) were used to predict the folding of the entire Gemin5 protein and to determine new possible construct targets. The gene fragment covering the predicted middle domain (residues 807–1097) was amplified by PCR using Phusion High-Fidelity DNA Polymerase (New England Biolabs), using pcDNA3-Xpress-G5 as template (Francisco-Velilla et al., 2016). Specific reverse and forward oligonucleotides (Sigma) are detailed in Table S1. Using the In-Fusion technology (Clontech), the gene fragment was inserted into the pOPINM vector (Oxford Protein Production Facility) that contains a MAH_6_SSG_MBP tag followed by a region cleavable by the PreScission protease.

The sequence encoding the p85 polypeptide present in pcDNA3-Xpress-G5 (Fernandez-Chamorro et al., 2014) was amplified by PCR with specific oligonucleotides (Table S1) and transferred to pcDNA3-CTAP (Chen and Gingras, 2007) via NotI-PacI to generate the construct pcDNA3-CTAP-p85. Constructs pcDNA3-CTAP-p85A951E and pcDNA3-Xpress-p85A951E were generated by QuickChange mutagenesis (Agilent Technologies) using specific primers (Table S1). Construct pOPINM_TPR-A951E was performed using PCR on pcDNA3-Xpress-p85A951E and In-Fusion cloning. Construct pcDNA3-Xpress-p85ΔRBS1/2 was generated by PCR amplifying the sequence present in pcDNA3-Xpress-G5 with specific primers (Table S1) and transferred to pcDNA3-Xpress via BamHI/NotI. All plasmids were confirmed by DNA sequencing (Macrogen). The construct expressing RBS1 protein was previously described (Francisco-Velilla et al., 2018).

### Expression and purification of G5-TPR

BL21 Rosetta(DE3)pLys cells transformed with the pOPINM_G5-TPR plasmid or the mutated variant pOPINM_G5-TPR-A951E were grown at 37°C in LB petri dishes supplemented with 2% glucose, 35 µg/ml ampicillin and 15 µg/ml chloramphenicol. Protein expression was performed in LB medium supplemented with 100 µg ml^−1^ ampicillin and 34 µg ml^−1^ chloramphenicol to mid-exponential phase (OD_600_ = 0.6–0.8). Protein expression was induced by adding 0.5 mM isopropyl-D-thiogalactopyranoside (IPTG) and incubating overnight at 20°C. Cells were harvested by centrifugation and stored at −80°C. The pellet from 1 L of culture was thawed and resuspended in 40 ml of buffer N_A_ (20 mM Tris-HCl pH 8.0, 0.5 M NaCl, 10 mM imidazole, 5% glycerol and 2 mM β-mercaptoethanol) supplemented with 1 pill of cOmplete^TM^ EDTA-free Protease Inhibitor cocktail (Sigma-Aldrich). Cells were then sonicated and the lysate was clarified by centrifugation in a Beckman JA-25.50 rotor at 50000 x g for 1 h at 4°C. The supernatant was filtered (0.45 µm pore) and loaded onto a 5 ml HisTrap FF column (GE Healthcare, USA). After extensively washing with buffer N_A_ supplemented with 35 mM imidazole, the protein was eluted by increasing the imidazole concentration to 250 mM in a single step. G5-TPR was cleaved overnight with GST-tagged PreScission protease (in a relation of 1/20^th^ of the protein weight) in buffer S_A-50_ (20 mM Tris-HCl pH 6.8, 50 mM NaCl, 1 mM DTT and 5% glycerol). Cleaved G5-TPR was loaded onto a 5 ml HiTrap S HP column (GE Healthcare, USA). Whereas the MAH_6_SSG_MBP tag is recovered in the flow-through, the protein of interest is retained in the column and eluted by increasing the salt concentration at 150–200 mM NaCl. After concentration through an Amicon Ultra system with a 10 kDa cut-off membrane, G5-TPR was further purified by size-exclusion chromatography on a Superdex 200 Increase 10/300 column (GE Healthcare) pre-equilibrated in buffer S_A-50_. The purified protein was concentrated as before and directly used for further studies or supplemented with 20% glycerol, flash-frozen in liquid nitrogen and stored at −80°C. All purification steps were carried out at 4°C. Sample purity was evaluated by SDS-PAGE with Coomassie staining. Protein concentration was determined by Bradford assay.

### Gel-filtration coupled to multi-angle light-scattering (SEC-MALS) measurements

500 µl of purified G5-TPR at 3 mg ml^−1^ was fractionated by gel filtration on a Superdex 200 10/300 column equilibrated in buffer S_A_, using ÄKTA purifier at a flow rate of 0.5 ml min^−1^. The eluted sample was characterized by in-line measurement of the refractive index and multi-angle light scattering using Optilab T-rEX and DAWN 8^+^ instruments, respectively (Wyatt). Data were analyzed using the *ASTRA* 6 software (Wyatt) to obtain the molar mass.

### Crystallization

Initial crystallization screenings were performed at RT with drops of 0.7 µl protein solution at 4.8 mg ml^−1^ plus 0.7 µl reservoir solution equilibrated against 60 µl of reservoir solution from JCSG+, PACT, MBP suite (Qiagen) and Crystal Screen (Hampton Research) commercial screens. Initial hits were further optimized in MRC 48-well sitting-drop plates (Molecular Dimensions). Best-diffracting plate-shaped crystals appeared after 3–5 days in 200 mM Na/K Tartrate, 25% PEG 3350 and 100 mM Bis-Tris-Methane pH 8.5. Other plate crystals grew in 200 mM NaI, 25% PEG 3350 and 100 mM Bis-Tris-Methane pH 8.5. In both cases, cryo-protection was reached by directly soaking the crystals in a solution containing the mother liquor supplemented with 20% glycerol. Crystals were then fished with cryo-loops and flash-cooled in liquid nitrogen.

### Data collection and structure determination

X-Ray diffraction data were remotely collected at BL13-XALOC (ALBA synchrotron, Barcelona) using a Pilatus 6M detector. For each set, a total wedge of 180° of data was collected with 0.15° oscillation and 0.1 s exposure per frame. Data processing and scaling were performed with XDS (Kabsch, 2010). Taking advantage of the anomalous edge scattering of the iodine present in the one of the crystallization conditions, crystallographic phases were determined by SAD using a wavelength of λ = 2.03 Å, where f ” for iodide is 10.5 e^−^ (6.1 keV; http://skuld.bmsc.washington.edu/scatter/). SAD phases were calculated with AutoSol in Phenix (Adams et al., 2010), which identified 13 iodide sites. Initial model building using a poly-Alanine sequence was performed with Autobuild and refined by iterative cycles with Phenix and Coot (Emsley et al., 2010). Suitable quality of the final models was validated according to MolProbity (Chen et al., 2010). Analysis of the macromolecular interfaces was performed with PDBePISA (Krissinel and Henrick, 2007) and InterProSurf (Negi et al., 2007). Sequence alignment and surface conservation representation were performed with Treefam (family TF328886), Ensembl (Genetree ENSGT00620000088064) and Consurf (Ashkenazy et al., 2016). Electrostatic surface potential was calculated with APBS plug-in of PyMOL and figures were prepared with PyMOL.

### RNA-DNA electrophoretic mobility shift assay

RNA probes were uniformly labeled using α^32^P-CTP (500 Ci/mmol), T7 RNA polymerase (10 U), and linearized DNA (1 µg), as described (Francisco-Velilla et al., 2015). Constructs expressing RNAs corresponding to domain 5 of the FMDV IRES or its single stranded region, the long structured RNA and the short hairpin, have been described (Francisco-Velilla et al., 2018; Pineiro et al., 2013). RNA was purified through MicroSpin G-25 Columns (GE Healthcare), ethanol precipitated and resuspended in Tris 10 mM, pH 8, EDTA 1mM (TE) to a final concentration of 0.04 pmol/µl. ssDNA and dsDNA probes (Table S1) were 5’-end labeled using T4 polynucleotide kinase and γ^32^P-ATP (500 Ci/mmol). RNA integrity and DNA probes were analyzed in 6% acrylamide 7 M urea denaturing gel electrophoresis.

RNA and DNA-binding reactions were carried out as described (Francisco-Velilla et al., 2018) with small modifications. The reactions were carried out in 10 µl of RNA-binding buffer [40 mM Tris-HCl pH 7.5, 250 mM NaCl, 0.1% (w/v) β-mercaptoethanol] for 15 min at room temperature. Electrophoresis was performed in non-denaturing 6.0% (29:1) polyacrylamide gels at 4°C, run in TBE buffer.

### Protein complexes isolation by Tandem Affinity Purification (TAP)

HEK293 cells (4 x P100), grown in Dulbecco’s Modified Eagle Medium (DMEM), transfected with the plasmids expressing p85-wt-TAP or p85-A951E-TAP proteins, were harvested 24 h post-transfection. The complexes associated to the TAP-tagged constructs were purified as described (Francisco-Velilla et al., 2015). Briefly, the supernatant of the first IgG Sepharose purification was subsequently subjected to a second Calmodulin (Agilent Technologies) purification step. Purified proteins were precipitated with 10% trichloroacetic acid at 4°C overnight, pelleted at 14000 g for 15 min at 4°C, washed three times with 1 ml of acetone and, finally, dissolved in SDS-loading buffer. A small aliquot was analyzed on silver stained SDS-PAGE gels to visualize the purification of proteins associated to Gemin5 p85-wt-TAP or p85-A951E-TAP polypeptides.

### In-Gel Digestion and Mass Spectrometry Analysis

Two independent biological replicates were analyzed for p85-TAP, and p85-A951E-TAP. The samples obtained by TAP were resuspended in 50 µl of sample buffer, and applied onto a 10% SDS-PAGE gel. Then, run was stopped as soon as the front entered 3 mm into the resolving gel. The protein bands concentrated in the stacking/resolving gel interface were visualized by Coomassie staining, excised, cut into cubes (2×2 mm), and placed in 0.5 ml tubes. The gel pieces were destained in acetonitrile:water (ACN:H_2_O, 1:1), were reduced and alkylated (disulfide bonds from cysteinyl residues were reduced with 10 mM DTT for 1 h at 56°C, and thiol groups were alkylated with 50 mM iodoacetamide for 1 h at room temperature in darkness) and digested in situ with sequencing grade trypsin (Promega, Madison, WI), as described (Shevchenko et al., 1996). The gel pieces were shrunk using ACN, which was then pipetted out and the gel pieces were dried in a speedvac. The dried gel pieces were re-swollen in 50 mM ammonium bicarbonate pH 8.8 with 60 ng/µl trypsin at 5:1 protein:trypsin (w/w) ratio. The tubes were kept in ice for 2 h and incubated at 37°C for 12 h. Digestion was stopped by the addition of 1% TFA. Whole supernatants were dried down and then desalted onto OMIX Pipette tips C18 (Agilent Technologies) until the mass spectrometric analysis.

The desalted protein digest was dried, resuspended in 10 µl of 0.1% formic acid and analyzed by RP-LC-MS/MS in an Easy-nLC II system coupled to an ion trap LTQ-Orbitrap-Velos-Pro hybrid mass spectrometer (Thermo Scientific). The peptides were concentrated (on-line) by reverse phase chromatography using a 0.1 mm × 20 mm C18 RP precolumn (Thermo Scientific), and then separated using a 0.075mm x 250 mm C18 RP column (Thermo Scientific) operating at 0.3 µl/min. Peptides were eluted using a 120 min dual gradient from 5 to 25% solvent B in 90 min followed by gradient from 25 to 40% solvent B over 120 min (Solvent A: 0.1% formic acid in water, solvent B: 0.1% formic acid, 80% acetonitrile in water). ESI ionization was done using a Nano-bore emitters Stainless Steel ID 30 µm (Proxeon) interface. The Orbitrap resolution was set at 30.000. Peptides were detected in survey scans from 400 to 1600 amu (1 µscan), followed by twenty data dependent MS/MS scans (Top 20), using an isolation width of 2 u (in mass-to-charge ratio units), normalized collision energy of 35%, and dynamic exclusion applied during 30 seconds periods.

Peptide identification from raw data was carried out using PEAKS Studio X (Zhang et al., 2012) search engine (Bioinformatics Solutions Inc). Database search was performed against UniProt-Homo sapiens FASTA (decoy-fusion database). The following constraints were used for the searches: tryptic cleavage after Arg and Lys, up to two missed cleavage sites, and tolerances of 20 ppm for precursor ions and 0.6 Da for MS/MS fragment ions and the searches were performed allowing optional Met oxidation and Cys carbamidomethylation. False discovery rates (FDR) for peptide spectrum matches (PSM) were limited to 0.01. Only those proteins with at least two distinct peptides being discovered from LC/MS/MS analyses were considered reliably identified.

### GO analysis

Gene Ontology analyses were performed by the DAVID database (https://david.ncifcrf.gov) on the overlapping proteins identified. The significantly enriched biological processes (BP) were identified using as a cutoff criteria P value <10^−5^ and a gene count ≥3.

### Gemin5 polypeptides expression and luciferase activity assays

HEK293 cell monolayers (2×10^5^) were cotransfected with a plasmid expressing luciferase in cap-dependent or IRES-dependent manner (pCAP-luc, pIRES-luc) (Lozano et al., 2018), and a plasmid expressing Xpress-p85-wt, Xpress-p85-A951E, Xpress-p85ΔRBS1/2, or the corresponding empty vector side by side using lipofectamine LTX (Thermo Scientific).

Cell lysates were prepared 24 h post-transfection in 100 µl lysis buffer (50 mM Tris-HCl pH 7.8, 100 mM NaCl, 0.5% NP40). The protein concentration in the lysate was determined by Bradford assay. Equal amounts of protein were loaded in SDS-PAGE and processed for western blotting to determine the expression of the polypeptides. Luciferase activity (RLU)/µg of total protein was internally normalized to the value obtained with the empty vector performed side by side. Each experiment was repeated independently three times. Values represent the mean ± SEM.

### Immunodetection

Xpress-p85, Xpress-p85-A951E and Xpress-p85ΔRBS1/2 proteins were immunodetected using anti-Xpress (Invitrogen) antibodies, and p85-wt-TAP and p85-A951E-TAP proteins were detected with anti-CBP (Abcam). Immunodetection of tubulin (Sigma) was used as loading control. Secondary antibodies (Thermo Scientific) were used according to the manufacturer’s instructions. The signal detected was done in the linear range of the antibodies.

### Statistical analyses

Statistical analyses for luciferase activity assays were performed as follows. Each experiment was repeated independently three times. Values represent the estimated mean ± SEM. We computed P values for a difference in distribution between two samples with the unpaired two-tailed Student’s *t*-test. Differences were considered significant when P < 0.05. The resulting P values were graphically illustrated in figures with asterisks as described in figure legends.

## SUPPLEMENTARY INFORMATION

**Supplementary Figure S1.**
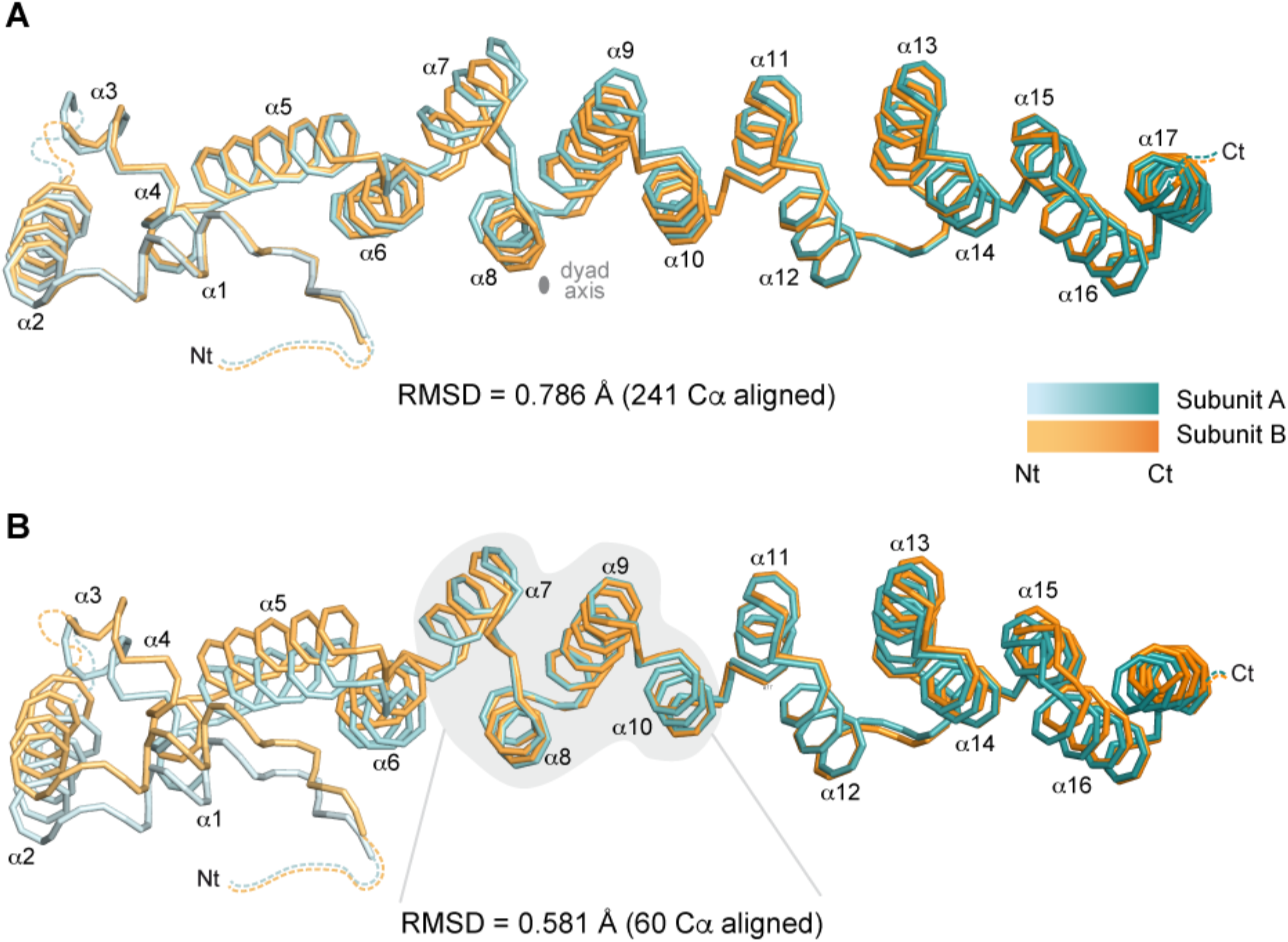
The two subunits in the G5-TPR dimer are similar. Related to Figure 1. Ribbon representation of the superposition between the two protein subunits within the crystal asymmetric unit, considering 241 Cα atoms **(A)**, or only helices α7–10 (60 Cα atoms; indicated in grey background) **(B)**. The value of the root-mean-square deviation (RMSD) for the superimposed atoms is indicated in each case. In **(A)** the position of the dyad axis is indicated.

**Supplementary Figure S2.**
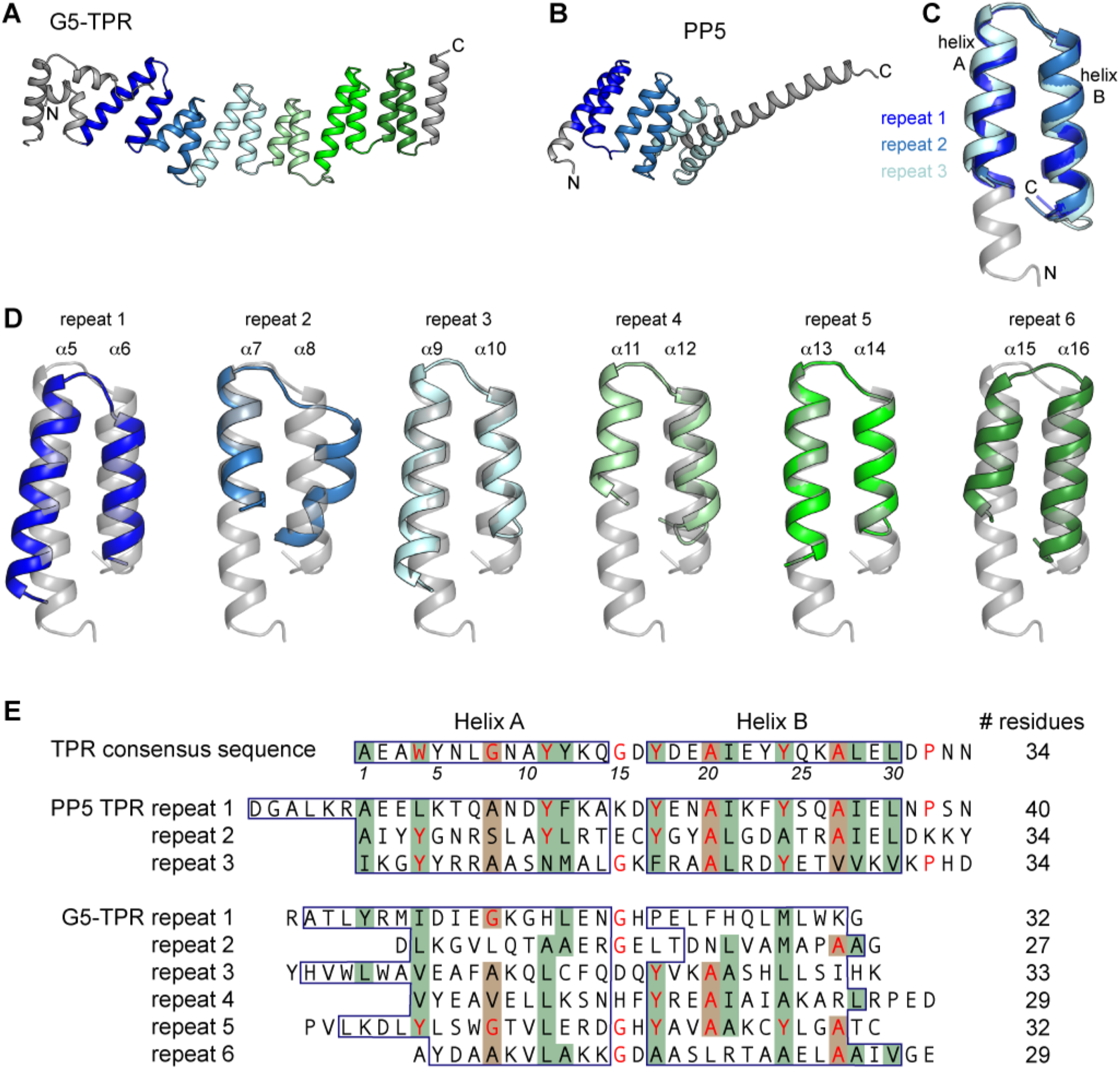
G5-TPR structure contains a tandem array of six TPR-like motifs. Related to Figure 1. **(A)** Crystal structure of one G5-TPR subunit with the six TPR-like helical pairs colored from blue to green. **(B)** Crystal structure of the TPR-domain of protein phosphatase PP5 (PDB: 1A17) (Das et al., 1998), with the three TPR motifs depicted in different colors and superimposed in **(C)**. **(D)** Superposition of the six helical repeats in G5-TPR with the first TPR motif of PP5 (colored in grey). **(E)** Sequence of the consensus TPR motif with highly conserved residues shown in red (D’Andrea and Regan, 2003), and sequence alignment of the three TPR motifs of PP5 and of the six TPR-like motifs of Gemin5, indicating the consensus pattern of small (green background) and large (brown background) hydrophobic residues, as defined in (Das et al., 1998).

**Supplementary Figure S3.**
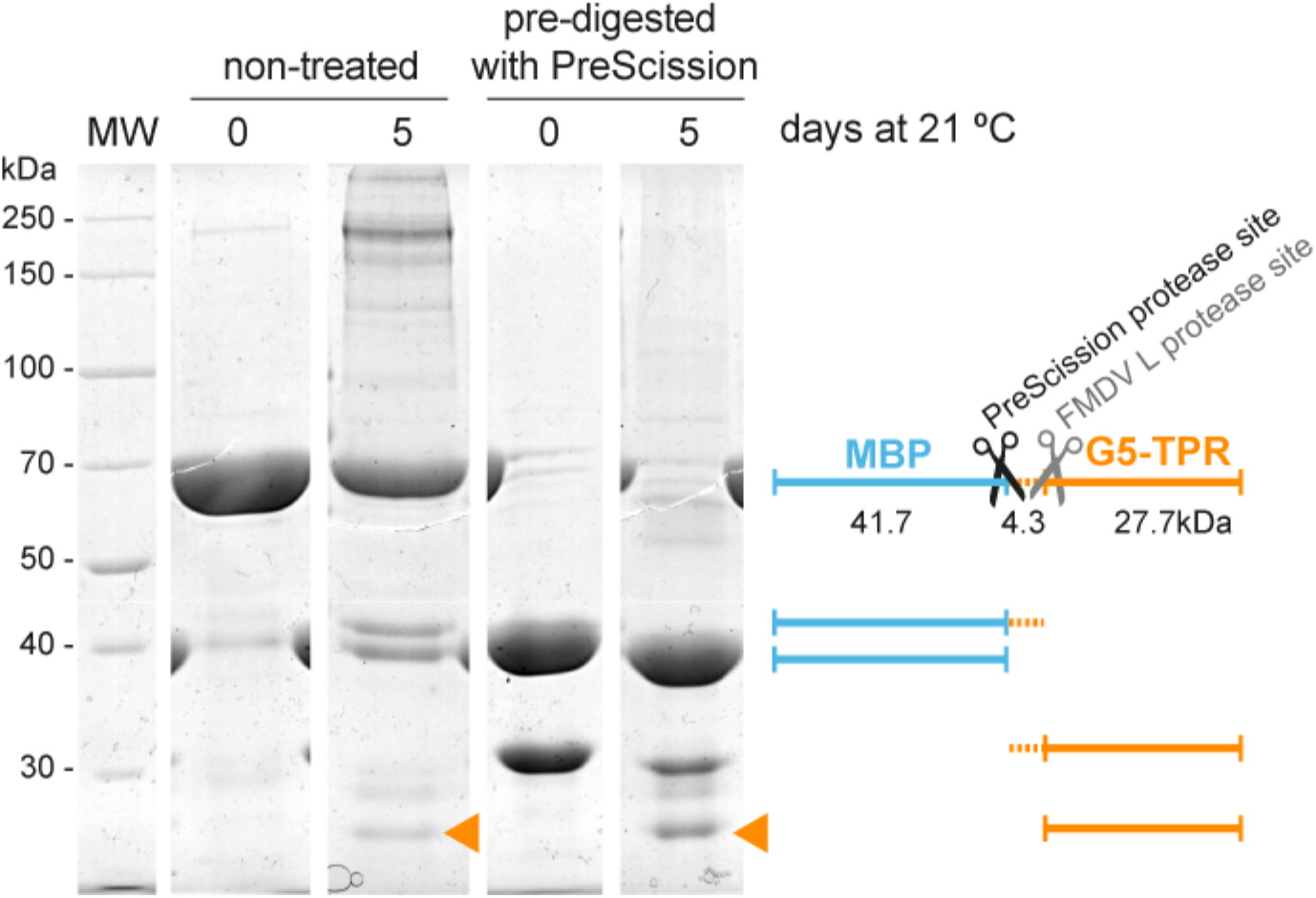
The Gemin5 TPR-like construct is prone to cleavage. Related to Figure 1. SDS-PAGE of partially purified G5-TPR samples non-treated or treated with the protease PreScission and incubated for 0 or 5 days at 21°C. The MBP-tagged G5-TPR construct has an expected molecular weight of 73.7 kDa. Bands of smaller size correspond to cleaved forms of the protein depicted on the right. The orange triangles indicate a protein fragment that matches the molecular weight of the G5-TPR crystal model.

**Supplementary Figure S4.**
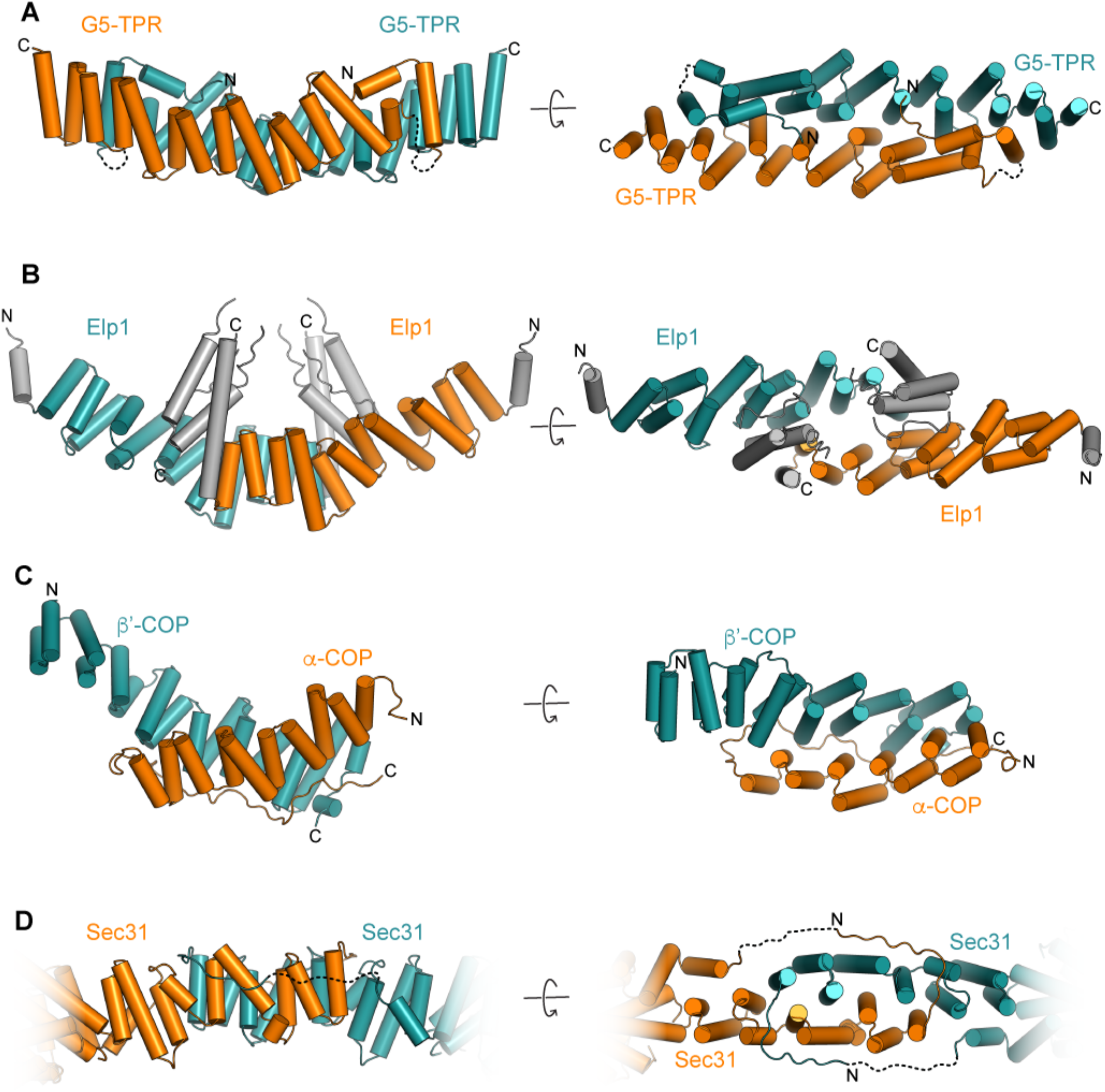
Structural similarity of G5-TPR dimer with other α-solenoid dimers. Related to Figure 1. (A, B) Cylinder representation of G5-TPR (A) and of human Elp1 TPR-like (PDB: 5CQR) (B) dimers in two perpendicular orientations. (C,D) Perpendicular views of the yeast vesicular coating β’-COP and α-COP heterodimer (PDB: 3MKQ) and of Sec31 homodimer (PDB: 2PM7).

**Supplementary Figure S5.**
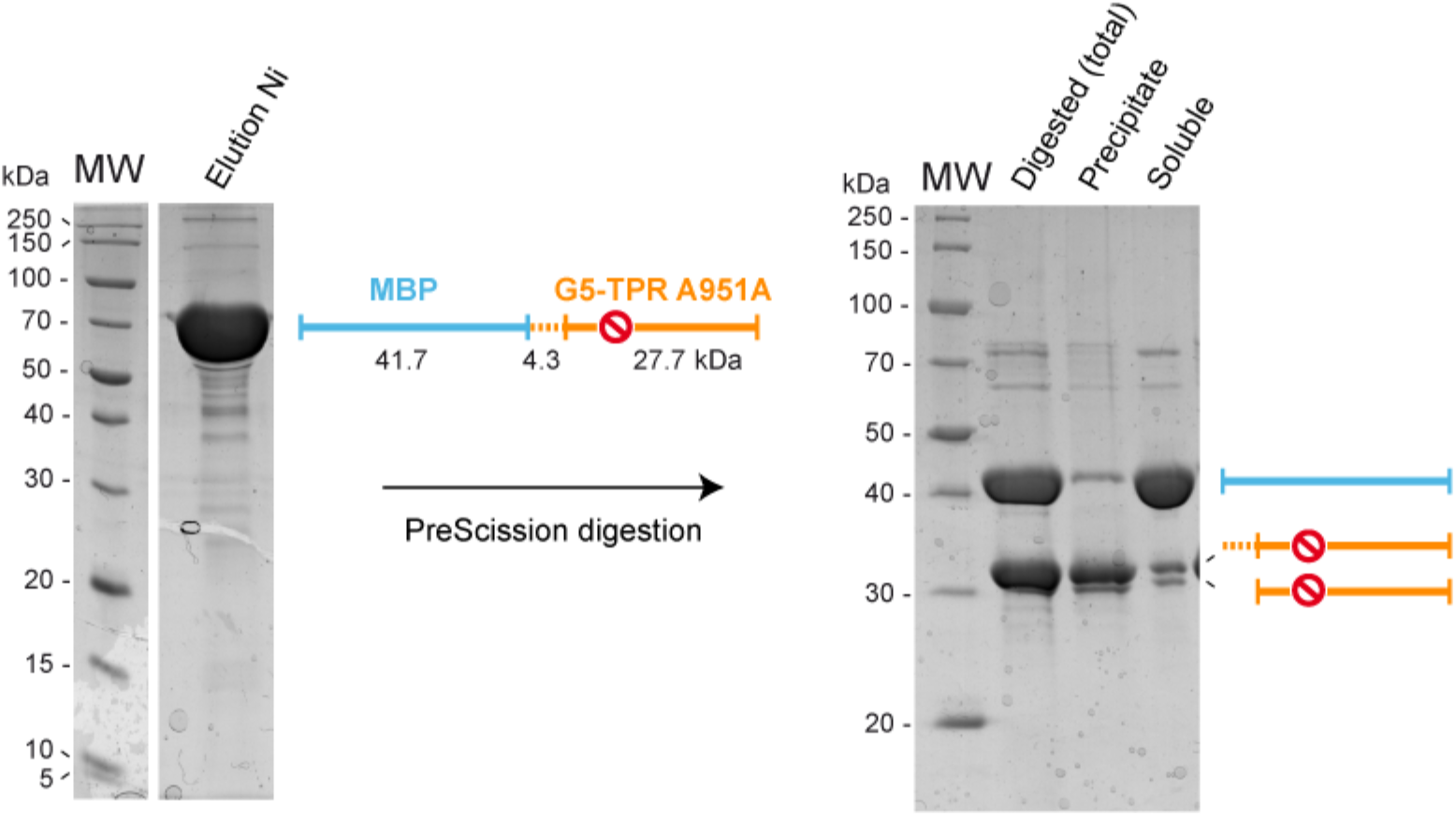
Mutant G5-TPR A951E is insoluble upon removal of the His_6_-MBP tag. Related to Figure. 1. Coomassie-stained SDS-PAGE of the His_6_-MBP tagged G5-TPR bearing mutation A951E, before (left gel) and after (right gel) digestion with PreScission protease, in a buffer solution containing 0.2 M NaCl. Upon removal of the His_6_-MBP tag the G5-TPR A951E protein is mostly in the insoluble fraction.

**Supplementary Figure S6.**
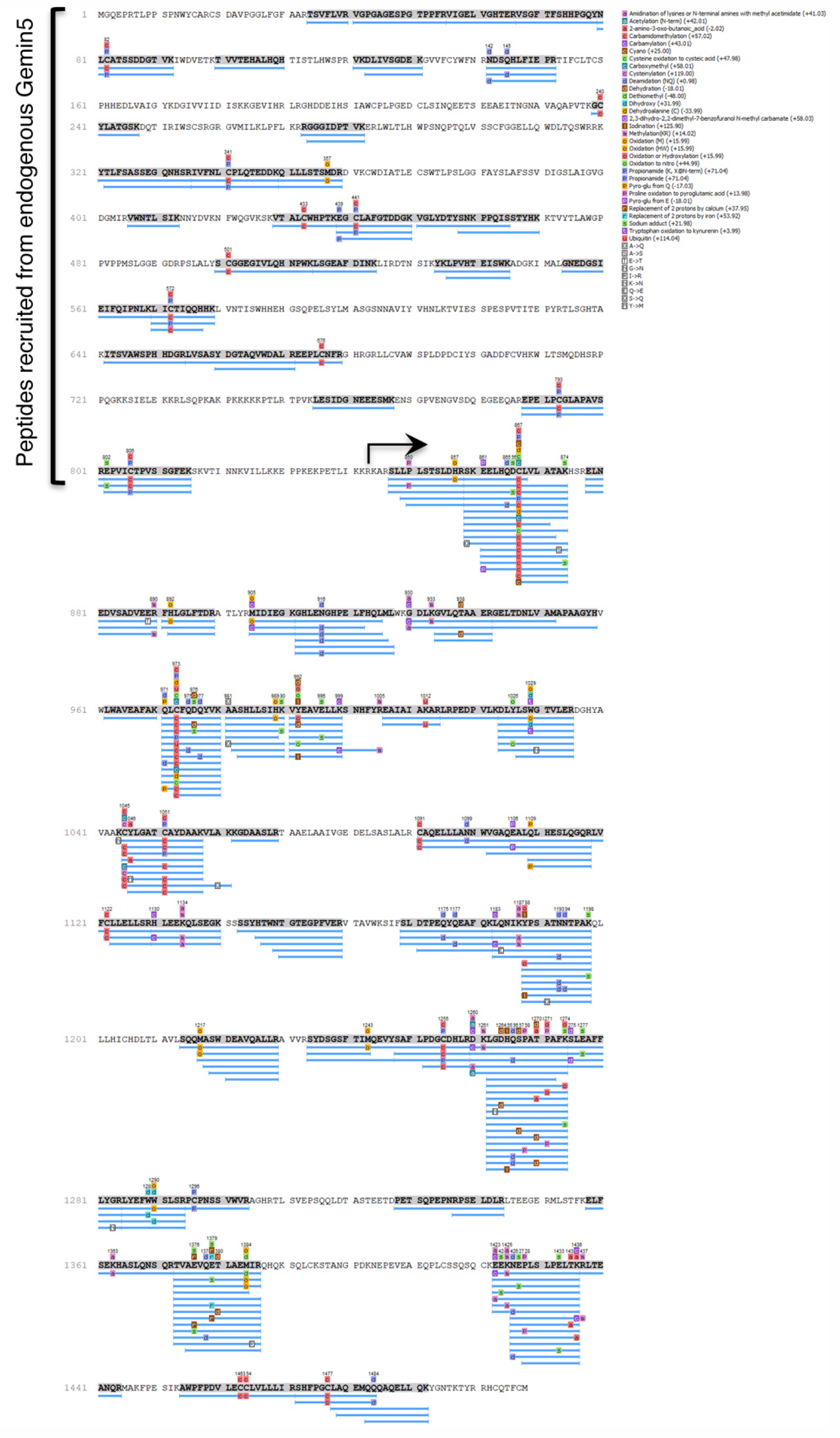
Peptide reads obtained in TAP pull-down assays carried out with p85-wt. Related to Figure 4. The Gemin5 peptides (blue lines) were identified from raw data with PEAKS. A bracket on the left depicts the amino acid region of the full-length Gemin5 absent in p85; an arrow denotes the first residue of the p85-wt protein fused to TAP tag expressed in HEK293 cells. The legend on the right depicts post-translational modifications.

**Supplementary Figure S7.**
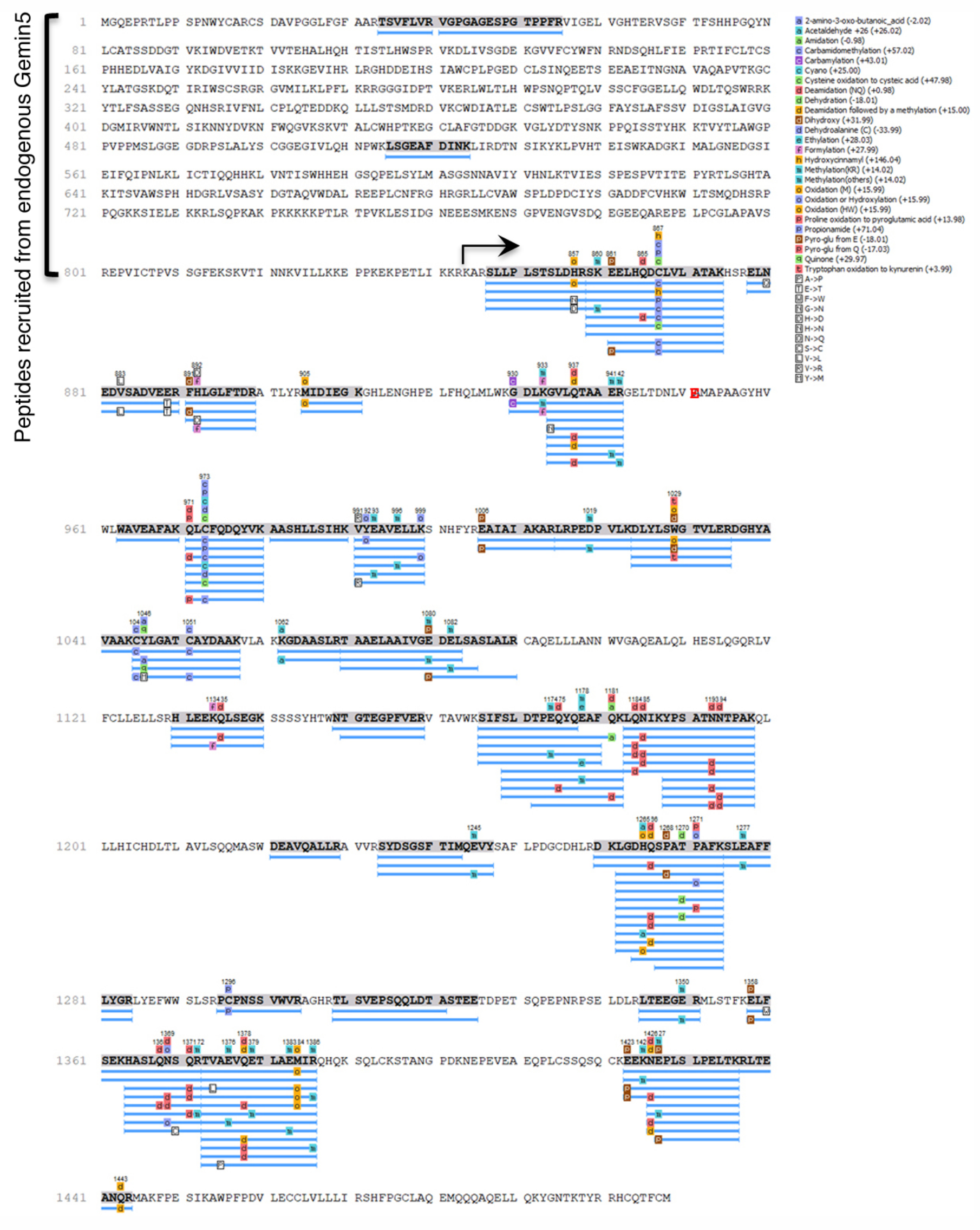
Peptide reads obtained in TAP pull-down assays carried out with p85-A951E. Related to Figure 4. The Gemin5 peptides (blue lines) were identified from raw data with PEAKS. A bracket on the left depicts the amino acid region absent in p85-A951E; the black arrow denotes the first residue of the p85-A951E protein fused to TAP tag expressed in HEK293 cells. The mutated residue A951E was verified by sequencing of the corresponding constructs; it was not detected in the MS/MS spectrum due to the large size of the trypsin fragment. The legend on the right depicts post-translational modifications.

**Table S1.**
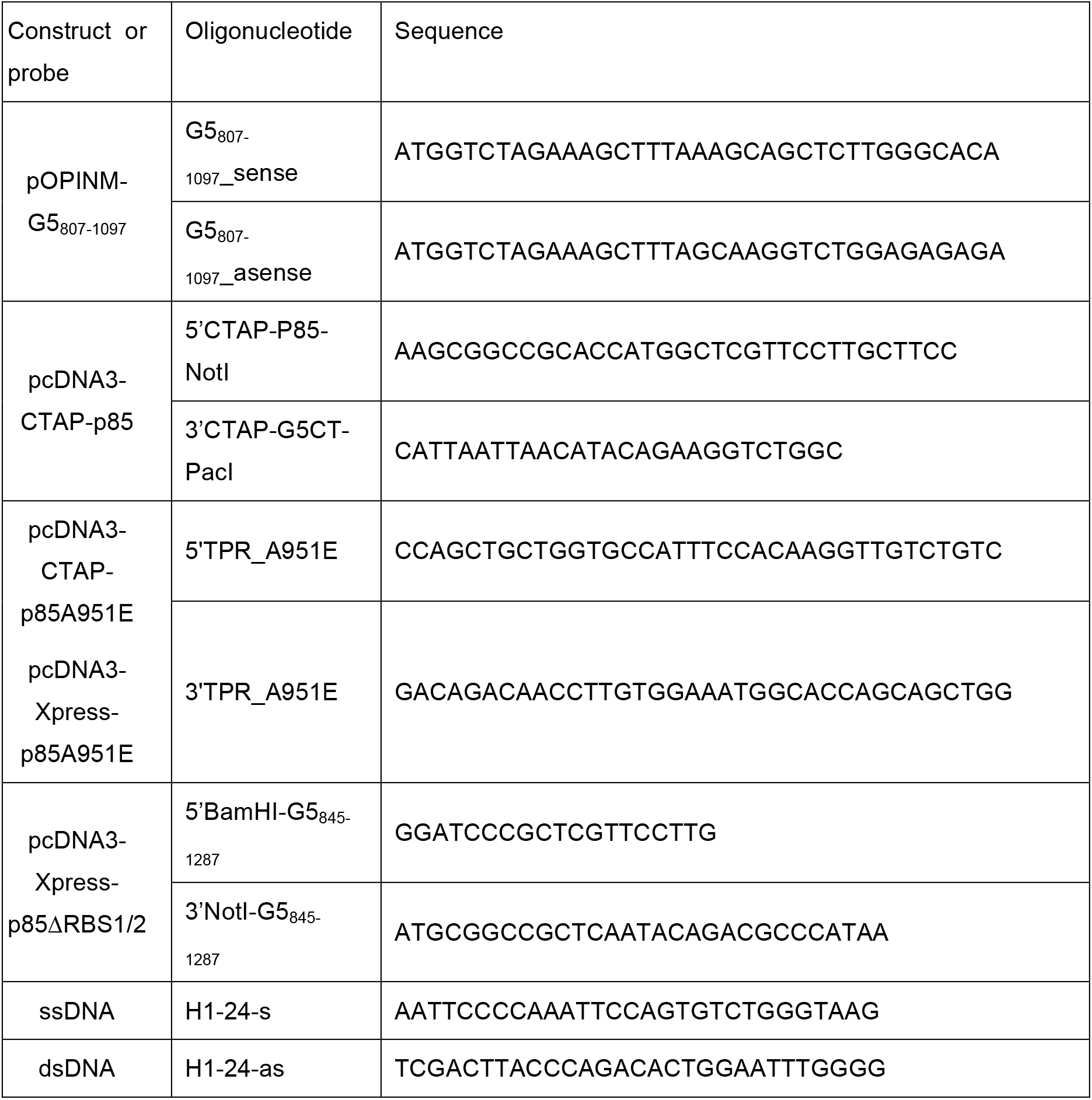
Oligonucleotides

## REFERENCES

Abbas, Y.M., Pichlmair, A., Gorna, M.W., Superti-Furga, G., and Nagar, B. (2013). Structural basis for viral 5’-PPP-RNA recognition by human IFIT proteins. Nature 494, 60–64.

Adams, P.D., Afonine, P.V., Bunkoczi, G., Chen, V.B., Davis, I.W., Echols, N., Headd, J.J., Hung, L.W., Kapral, G.J., Grosse-Kunstleve, R.W., et al. (2010). PHENIX: a comprehensive Python-based system for macromolecular structure solution. Acta Crystallogr D Biol Crystallogr 66, 213–221.

Ashkenazy, H., Abadi, S., Martz, E., Chay, O., Mayrose, I., Pupko, T., and Ben-Tal, N. (2016). ConSurf 2016: an improved methodology to estimate and visualize evolutionary conservation in macromolecules. Nucleic Acids Res 44, W344–350.

Battle, D.J., Kasim, M., Wang, J., and Dreyfuss, G. (2007). SMN-independent subunits of the SMN complex. Identification of a small nuclear ribonucleoprotein assembly intermediate. J Biol Chem 282, 27953–27959.

Battle, D.J., Lau, C.K., Wan, L., Deng, H., Lotti, F., and Dreyfuss, G. (2006). The Gemin5 protein of the SMN complex identifies snRNAs. Mol Cell 23, 273–279.

Burghes, A.H., and Beattie, C.E. (2009). Spinal muscular atrophy: why do low levels of survival motor neuron protein make motor neurons sick? Nat Rev Neurosci 10, 597–609.

Chen, G.I., and Gingras, A.C. (2007). Affinity-purification mass spectrometry (AP-MS) of serine/threonine phosphatases. Methods 42, 298–305.

Chen, V.B., Arendall, W.B., 3rd, Headd, J.J., Keedy, D.A., Immormino, R.M., Kapral, G.J., Murray, L.W., Richardson, J.S., and Richardson, D.C. (2010). MolProbity: all-atom structure validation for macromolecular crystallography. Acta Crystallogr D Biol Crystallogr 66, 12–21.

D’Andrea, L.D., and Regan, L. (2003). TPR proteins: the versatile helix. Trends Biochem Sci 28, 655–662.

Das, A.K., Cohen, P.W., and Barford, D. (1998). The structure of the tetratricopeptide repeats of protein phosphatase 5: implications for TPR-mediated protein-protein interactions. EMBO J 17, 1192–1199.

Ellisdon, A.M., Dimitrova, L., Hurt, E., and Stewart, M. (2012). Structural basis for the assembly and nucleic acid binding of the TREX-2 transcription-export complex. Nat Struct Mol Biol 19, 328–336.

Emsley, P., Lohkamp, B., Scott, W.G., and Cowtan, K. (2010). Features and development of Coot. Acta Crystallogr D Biol Crystallogr 66, 486–501.

Fath, S., Mancias, J.D., Bi, X., and Goldberg, J. (2007). Structure and organization of coat proteins in the COPII cage. Cell 129, 1325–1336.

Fernandez-Chamorro, J., Pineiro, D., Gordon, J.M., Ramajo, J., Francisco-Velilla, R., Macias, M.J., and Martinez-Salas, E. (2014). Identification of novel non-canonical RNA-binding sites in Gemin5 involved in internal initiation of translation. Nucleic Acids Res 42, 5742–5754.

Francisco-Velilla, R., Azman, E.B., and Martinez-Salas, E. (2019). Impact of RNA-Protein Interaction Modes on Translation Control: The Versatile Multidomain Protein Gemin5. Bioessays 41, e1800241.

Francisco-Velilla, R., Fernandez-Chamorro, J., Dotu, I., and Martinez-Salas, E. (2018). The landscape of the non-canonical RNA-binding site of Gemin5 unveils a feedback loop counteracting the negative effect on translation. Nucleic Acids Res 46, 7339–7353.

Francisco-Velilla, R., Fernandez-Chamorro, J., Lozano, G., Diaz-Toledano, R., and Martinez-Salas, E. (2015). RNA-protein interaction methods to study viral IRES elements. Methods 91, 3–12.

Francisco-Velilla, R., Fernandez-Chamorro, J., Ramajo, J., and Martinez-Salas, E. (2016). The RNA-binding protein Gemin5 binds directly to the ribosome and regulates global translation. Nucleic Acids Res 44, 8335–8351.

Garcia-Moreno, M., Noerenberg, M., Ni, S., Jarvelin, A.I., Gonzalez-Almela, E., Lenz, C.E., Bach-Pages, M., Cox, V., Avolio, R., Davis, T., et al. (2019). System-wide Profiling of RNA-Binding Proteins Uncovers Key Regulators of Virus Infection. Mol Cell 74, 196–211.e11.

Gates, J., Lam, G., Ortiz, J.A., Losson, R., and Thummel, C.S. (2004). rigor mortis encodes a novel nuclear receptor interacting protein required for ecdysone signaling during Drosophila larval development. Development 131, 25–36.

Gehring, N.H., Wahle, E., and Fischer, U. (2017). Deciphering the mRNP Code: RNA-Bound Determinants of Post-Transcriptional Gene Regulation. Trends Biochem Sci 42, 369–382.

Grotwinkel, J.T., Wild, K., Segnitz, B., and Sinning, I. (2014). SRP RNA remodeling by SRP68 explains its role in protein translocation. Science 344, 101–104.

Gundogdu, M., Llabres, S., Gorelik, A., Ferenbach, A.T., Zachariae, U., and van Aalten, D.M.F. (2018). The O-GlcNAc Transferase Intellectual Disability Mutation L254F Distorts the TPR Helix. Cell Chem Biol 25, 513–518 e514.

Holm, L., and Sander, C. (1995). Dali: a network tool for protein structure comparison. Trends Biochem Sci 20, 478–480.

Huang da, W., Sherman, B.T., and Lempicki, R.A. (2009). Systematic and integrative analysis of large gene lists using DAVID bioinformatics resources. Nat Protoc 4, 44–57.

Jin, W., Wang, Y., Liu, C.P., Yang, N., Jin, M., Cong, Y., Wang, M., and Xu, R.M. (2016). Structural basis for snRNA recognition by the double-WD40 repeat domain of Gemin5. Genes Dev 30, 2391–2403.

Johnson, B., VanBlargan, L.A., Xu, W., White, J.P., Shan, C., Shi, P.Y., Zhang, R., Adhikari, J., Gross, M.L., Leung, D.W., et al. (2018). Human IFIT3 Modulates IFIT1 RNA Binding Specificity and Protein Stability. Immunity 48, 487–499 e485.

Jonas, S., and Izaurralde, E. (2013). The role of disordered protein regions in the assembly of decapping complexes and RNP granules. Genes Dev 27, 2628–2641.

Kabsch, W. (2010). Xds. Acta Crystallogr D Biol Crystallogr 66, 125–132.

Kajander, T., Cortajarena, A.L., Mochrie, S., and Regan, L. (2007). Structure and stability of designed TPR protein superhelices: unusual crystal packing and implications for natural TPR proteins. Acta Crystallogr D Biol Crystallogr 63, 800–811.

Katibah, G.E., Lee, H.J., Huizar, J.P., Vogan, J.M., Alber, T., and Collins, K. (2013). tRNA binding, structure, and localization of the human interferon-induced protein IFIT5. Mol Cell 49, 743–750.

Kelley, L.A., Mezulis, S., Yates, C.M., Wass, M.N., and Sternberg, M.J. (2015). The Phyre2 web portal for protein modeling, prediction and analysis. Nat Protoc 10, 845–858.

Kim, M.S., Pinto, S.M., Getnet, D., Nirujogi, R.S., Manda, S.S., Chaerkady, R., Madugundu, A.K., Kelkar, D.S., Isserlin, R., Jain, S., et al. (2014). A draft map of the human proteome. Nature 509, 575–581.

Krissinel, E., and Henrick, K. (2007). Inference of macromolecular assemblies from crystalline state. J Mol Biol 372, 774–797.

Lau, C.K., Bachorik, J.L., and Dreyfuss, G. (2009). Gemin5-snRNA interaction reveals an RNA binding function for WD repeat domains. Nat Struct Mol Biol 16, 486–491.

Lee, C., and Goldberg, J. (2010). Structure of coatomer cage proteins and the relationship among COPI, COPII, and clathrin vesicle coats. Cell 142, 123–132.

Lozano, G., Francisco-Velilla, R., and Martinez-Salas, E. (2018). Ribosome-dependent conformational flexibility changes and RNA dynamics of IRES domains revealed by differential SHAPE. Sci Rep 8, 5545.

Lozano, G., and Martinez-Salas, E. (2015). Structural insights into viral IRES-dependent translation mechanisms. Curr Opin Virol 12, 113–120.

Lunde, B.M., Moore, C., and Varani, G. (2007). RNA-binding proteins: modular design for efficient function. Nat Rev Mol Cell Biol 8, 479–490.

Matera, A.G., Raimer, A.C., Schmidt, C.A., Kelly, J.A., Droby, G.N., Baillat, D., Ten Have, S., Lamond, A.I., Wagner, E.J., and Gray, K.M. (2019). Composition of the Survival Motor Neuron (SMN) Complex in Drosophila melanogaster. G3 (Bethesda) 9, 491–503.

Negi, S.S., Schein, C.H., Oezguen, N., Power, T.D., and Braun, W. (2007). InterProSurf: a web server for predicting interacting sites on protein surfaces. Bioinformatics 23, 3397–3399.

Pacheco, A., Lopez de Quinto, S., Ramajo, J., Fernandez, N., and Martinez-Salas, E. (2009). A novel role for Gemin5 in mRNA translation. Nucleic Acids Res 37, 582–590.

Piazzon, N., Schlotter, F., Lefebvre, S., Dodre, M., Mereau, A., Soret, J., Besse, A., Barkats, M., Bordonne, R., Branlant, C., et al. (2013). Implication of the SMN complex in the biogenesis and steady state level of the signal recognition particle. Nucleic Acids Res 41, 1255–1272.

Pineiro, D., Fernandez, N., Ramajo, J., and Martinez-Salas, E. (2013). Gemin5 promotes IRES interaction and translation control through its C-terminal region. Nucleic Acids Res 41, 1017–1028.

Pineiro, D., Ramajo, J., Bradrick, S.S., and Martinez-Salas, E. (2012). Gemin5 proteolysis reveals a novel motif to identify L protease targets. Nucleic Acids Res 40, 4942–4953.

Sacristan-Reviriego, A., Bellingham, J., Prodromou, C., Boehm, A.N., Aichem, A., Kumaran, N., Bainbridge, J., Michaelides, M., and van der Spuy, J. (2017). The integrity and organization of the human AIPL1 functional domains is critical for its role as a HSP90-dependent co-chaperone for rod PDE6. Hum Mol Genet 26, 4465–4480.

Shevchenko, A., Wilm, M., Vorm, O., and Mann, M. (1996). Mass spectrometric sequencing of proteins silver-stained polyacrylamide gels. Anal Chem 68, 850–858.

Simsek, D., Tiu, G.C., Flynn, R.A., Byeon, G.W., Leppek, K., Xu, A.F., Chang, H.Y., and Barna, M. (2017). The Mammalian Ribo-interactome Reveals Ribosome Functional Diversity and Heterogeneity. Cell 169, 1051–1065 e1018.

Tang, X., Bharath, S.R., Piao, S., Tan, V.Q., Bowler, M.W., and Song, H. (2016). Structural basis for specific recognition of pre-snRNA by Gemin5. Cell Res 26, 1353–1356.

Uhlen, M., Fagerberg, L., Hallstrom, B.M., Lindskog, C., Oksvold, P., Mardinoglu, A., Sivertsson, A., Kampf, C., Sjostedt, E., Asplund, A., et al. (2015). Proteomics. Tissue-based map of the human proteome. Science 347, 1260419.

Walsh, D., and Mohr, I. (2011). Viral subversion of the host protein synthesis machinery. Nat Rev Microbiol 9, 860–875.

Workman, E., Kalda, C., Patel, A., and Battle, D.J. (2015). Gemin5 Binds to the Survival Motor Neuron mRNA to Regulate SMN Expression. J Biol Chem, 290, 15662–15669.

Xu, C., Ishikawa, H., Izumikawa, K., Li, L., He, H., Nobe, Y., Yamauchi, Y., Shahjee, H.M., Wu, X.H., Yu, Y.T., et al. (2016). Structural insights into Gemin5-guided selection of pre-snRNAs for snRNP assembly. Genes Dev 30, 2376–2390.

Xu, H., Lin, Z., Li, F., Diao, W., Dong, C., Zhou, H., Xie, X., Wang, Z., Shen, Y., and Long, J. (2015). Dimerization of elongator protein 1 is essential for Elongator complex assembly. Proc Nat Acad Sci U S A 112, 10697–10702.

Yang, J., Yan, R., Roy, A., Xu, D., Poisson, J., and Zhang, Y. (2015). The I-TASSER Suite: protein structure and function prediction. Nat Methods 12, 7–8.

Yang, Z., Liang, H., Zhou, Q., Li, Y., Chen, H., Ye, W., Chen, D., Fleming, J., Shu, H., and Liu, Y. (2012). Crystal structure of ISG54 reveals a novel RNA binding structure and potential functional mechanisms. Cell Res 22, 1328–1338.

Yong, J., Kasim, M., Bachorik, J.L., Wan, L., and Dreyfuss, G. (2010). Gemin5 delivers snRNA precursors to the SMN complex for snRNP biogenesis. Mol Cell 38, 551–562.

Zhang, J., Xin, L., Shan, B., Chen, W., Xie, M., Yuen, D., Zhang, W., Zhang, Z., Lajoie, G.A., and Ma, B. (2012). PEAKS DB: de novo sequencing assisted database search for sensitive and accurate peptide identification. Mol Cell Prot : MCP 11, M111 010587.

Zhou, X., Liao, H., Chern, M., Yin, J., Chen, Y., Wang, J., Zhu, X., Chen, Z., Yuan, C., Zhao, W., et al. (2018). Loss of function of a rice TPR-domain RNA-binding protein confers broad-spectrum disease resistance. Proc Nat Acad Sci U S A 115, 3174–3179.

Zhu, H., Sepulveda, E., Hartmann, M.D., Kogenaru, M., Ursinus, A., Sulz, E., Albrecht, R., Coles, M., Martin, J., and Lupas, A.N. (2016). Origin of a folded repeat protein from an intrinsically disordered ancestor. Elife 5.

Zimmermann, L., Stephens, A., Nam, S.Z., Rau, D., Kubler, J., Lozajic, M., Gabler, F., Soding, J., Lupas, A.N., and Alva, V. (2018). A Completely Reimplemented MPI Bioinformatics Toolkit with a New HHpred Server at its Core. J Mol Biol 430, 2237–2243.

